# Brilacidin, a novel antifungal agent against *Cryptococcus neoformans*

**DOI:** 10.1101/2024.04.10.588976

**Authors:** Camila Diehl, Camila Figueiredo Pinzan, Patrícia Alves de Castro, Endrews Delbaje, Laura C. García Carnero, Eddy Sánchez-León, Kabir Bhalla, James W. Kronstad, Dong-gyu Kim, Tamara L. Doering, Sondus Alkhazraji, Nagendra N. Mishra, Ashraf S. Ibrahim, Mami Yoshimura, Luis Alberto Vega Isuhuaylas, Lien Thi Kim Pham, Yoko Yashiroda, Charles Boone, Thaila Fernanda dos Reis, Gustavo H. Goldman

## Abstract

*Cryptococcus neoformans* causes cryptococcosis, one of the most prevalent fungal diseases, generally characterized by meningitis. There is a limited and not very effective number of drugs available to combat this disease. In this manuscript, we show the host defense peptide mimetic brilacidin (BRI) as a promising antifungal drug against *C. neoformans*. BRI is able to affect the organization of the cell membrane, increasing fungal cell permeability. We also investigated the effects of BRI against the model system *Saccharomyces cerevisiae* by analyzing libraries of mutants grown in the presence of BRI. In *S. cerevisiae*, BRI also affects the cell membrane organization, but in addition the cell wall integrity pathway and calcium metabolism. *In vivo* experiments show BRI significantly reduces *C. neoformans* survival inside macrophages and partially clears *C. neoformans* lung infection in an immunocompetent murine model of invasive pulmonary cryptococcosis. We also observed that BRI interacts with caspofungin (CAS) and amphotericin (AmB), potentiating their mechanism of action against *C. neoformans*. BRI+CAS affects endocytic movement, calcineurin, and mitogen activated protein kinases. Our results indicate that BRI is a novel antifungal drug against cryptococcosis.

**Importance:** Invasive fungal infections have a high mortality rate causing more deaths annually than tuberculosis or malaria. Cryptococcosis, one of the most prevalent fungal diseases, is generally characterized by meningitis and is mainly caused by two closely related species of basidiomycetous yeasts, *Cryptococcus neoformans* and *Cryptococcus gattii*. There are few therapeutic options for treating cryptococcosis and searching for new antifungal agents against this disease is very important. Here, we present brilacidin (BRI) as a potential antifungal agent against *C. neoformans*. BRI is a small molecule host defense peptide mimetic that has previously exhibited broad-spectrum immunomodulatory/anti-inflammatory activity against bacteria and viruses. BRI alone was shown to inhibit the growth of *C. neoformans*, acting as a fungicidal drug, but surprisingly also potentiated the activity of caspofungin (CAS) against this species. We investigated the mechanism of action of BRI and BRI+CAS against *C. neoformans*. We propose BRI as a new antifungal agent against cryptococcosis.

## Introduction

Fungal pathogens are a major global health problem affecting more than a billion people worldwide^1,2^. Invasive fungal infections have a high mortality rate causing more deaths annually than tuberculosis or malaria^3^. One of the most prevalent fungal diseases, cryptococcosis, is generally characterized by meningitis and is mainly caused by two closely related species of basidiomycetous yeasts, *Cryptococcus neoformans* and *Cryptococcus gattii*^4,5^. These two pathogenic species share well-characterized virulence determinants, such as the presence of a polysaccharide capsule, the synthesis of melanin, the ability to proliferate at human body temperature and inside of macrophages, tolerance to CO_2_, and the production of host-damaging enzymes like urease and phospholipase^6–10^. These traits contribute to their survival and success as human pathogens^11,12^.

The global incidence of cryptococcal meningitis is estimated at more than 400,000 new cases annually, with 181,100 annual deaths^13,14^. This number corresponds to approximately 19% of AIDS-related deaths globally^14,15^. Despite the significant impact of these infections, treatment options for cryptococcosis remain limited^16,17^. The polyenes, azoles and echinocandins are the three major classes of drugs used for treatment of invasive fungal infections^18^. Polyenes and azoles target ergosterol, depleting the lipid at the fungal plasma membrane or directly blocking its biosynthesis, while the echinocandins act by inhibiting the production of (1,3)-β-D-glucan to disrupt fungal cell wall integrity^16,19^. Amphotericin B (AmB), an antifungal of the polyene class, is the first option for treatment of cryptococcal infections, but it is hampered by substantial toxicity and high cost^20,21^. The success of a combination regimen of amphotericin B plus flucytosine (a compound that inhibits pyrimidine biosynthesis;^22^) led to updated guidelines for the treatment of cryptococcal disease in HIV-patients^20^. However, in low- and middle-income countries the scarcity of these two drugs has resulted in widespread use of much less effective fluconazole monotherapy, which explains the dramatically high mortality rate^23^.

Cryptococcal cells are intrinsically resistant to echinocandins and they employ an arsenal of mechanisms that enable resistance to azoles, such as upregulation of the azole target gene *ERG11* ^24,25^. One is the genomic plasticity that enables cells to form heteroresistant strains under azole stress^26,27^. This heteroresistance can be intrinsic and present in all strains regardless of prior drug exposure^28^. Genomic plasticity can further enable resistance through the movement of transposable elements, which have been recently linked to drug resistance in *C. neoformans*^29^. Together, these genomic changes allow *Cryptococcus* spp. a great capacity for physiological adaptation, such that they respond quickly and efficiently to stress in the environment and frequently evolve drug resistance^30,31^. Other features of *C. neoformans*, including a complex cell wall and polysaccharide capsule and the ability to undergo dramatic morphological transitions, contribute to both virulence and drug resistance^32–34^. The plasma membrane also plays an important role in antifungal resistance^35^. For example, transporters present in the cell membrane may act as antifungal efflux pumps contributing to azole resistance in *C. neoformans* and *C. gattii*^36^. Maintenance of lipid asymmetry of the phospholipid membrane is also involved in caspofungin (CAS) resistance^37–39^.

Taking into consideration the increased number of individuals susceptible to cryptococcal infection, the search for new antifungal agents has become more relevant than ever. Here, we present brilacidin (BRI) as a potential antifungal agent against *C. neoformans*. BRI is a small molecule host defense peptide mimetic that has previously exhibited broad-spectrum immunomodulatory/anti-inflammatory activity against bacteria and viruses^40^. Recently, we showed that BRI potentiates the effect of azoles and CAS against several fungal species, decreasing *Aspergillus fumigatus* fungal burden in a murine chemotherapeutic model of invasive pulmonary aspergillosis and ablating disease development in a murine model of fungal keratitis^41^. BRI alone was shown to inhibit the growth of *C. neoformans*, acting as a fungicidal drug at concentrations of 2.5 µM, but surprisingly also potentiated the activity of CAS against this species.

Here, we investigated the mechanism of action (MoA) of BRI and BRI+CAS against *C. neoformans*. We observed that not only *C. neoformans* but also *C. gattii* and several clinical isolates from both species are susceptible to BRI, and the combination of BRI+CAS has an additive effect against these strains. We screened a library of *C. neoformans* transcription factor mutants and found that deletion of the gene that encodes the sterol regulatory element-binding 1 protein (*SRE1*) yielded cells showing significant sensitivity to BRI alone. Sre1 is a transcriptional activator required for adaptation to hypoxic growth, which plays an essential role in the ergosterol biosynthesis pathway^42^. *C. neoformans* mutants defective in ergosterol and glycosphingolipid synthesis have reduced MIC values for BRI and supplementation with free ergosterol reverses the inhibitory effects of BRI, suggesting that this compound affects cell membrane organization. We implicated casein kinases in the BRI MoA, likely via modulating the expression of genes involved in ergosterol biosynthesis and cell wall construction. Analysis of a library of *S. cerevisiae* mutants grown in the presence of BRI indicated that fungal cell membrane organization, cell wall integrity, and calcium metabolism are affected by this compound. *C. neoformans* endocytic machinery plays a role in the MoA of BRI+CAS combinations, affecting cell permeability, but the Cell Wall Integrity (CWI) pathway is also important for this MoA, affecting mainly chitin accumulation in the cell wall. Finally, we showed that BRI significantly reduces *C. neoformans* survival inside macrophages and partially clears *C. neoformans* lung infection in an immunocompetent murine model of invasive pulmonary cryptococcosis. We propose BRI as a new antifungal agent against cryptococcosis.

## Results

### The MoA of BRI is dependent on the disorganization of the cell membrane in *C. neoformans*

Previously, we have shown that BRI can fungicidally inhibit *C. neoformans* H99 growth^41^. Not only is this strain susceptible to BRI, but *C. neoformans* KN99a and *C. gattii* R265, and six clinical isolates, three from each species are also sensitive to BRI (**Table 1**). *C. neoformans* clinical isolates were assayed for mutation rates by using a modified Luria-Delbruck fluctuation test^43^, for 5-fluorocytosine (5-FC) or BRI (**Figure 1a**). Replicate cultures grown without selection were challenged on RPMI medium containing 5-FC or BRI to determine the probability that cells would spontaneously gain mutations that provide resistance (**Figure 1a**). On 5-FC, strain H99 and clinical isolate 478 had a spontaneous mutation rate of 1.59 x 10^-8^ (95% confidence interval: 6.15 x 10^-9^ to 2.56 x 10^-8^) and 2.37 x 10^-9^ (95% confidence interval: 3.61 x 10^-10^ to 4.37 x 10^-9^), respectively, while no BRI-resistant mutants were observed on the RPMI medium containing BRI (**Figure 1a**). These results indicate that mutations that confer resistance to BRI were much less likely to occur spontaneously in *C. neoformans* cells than those that confer resistance to 5-FC.

**Figure 1.**
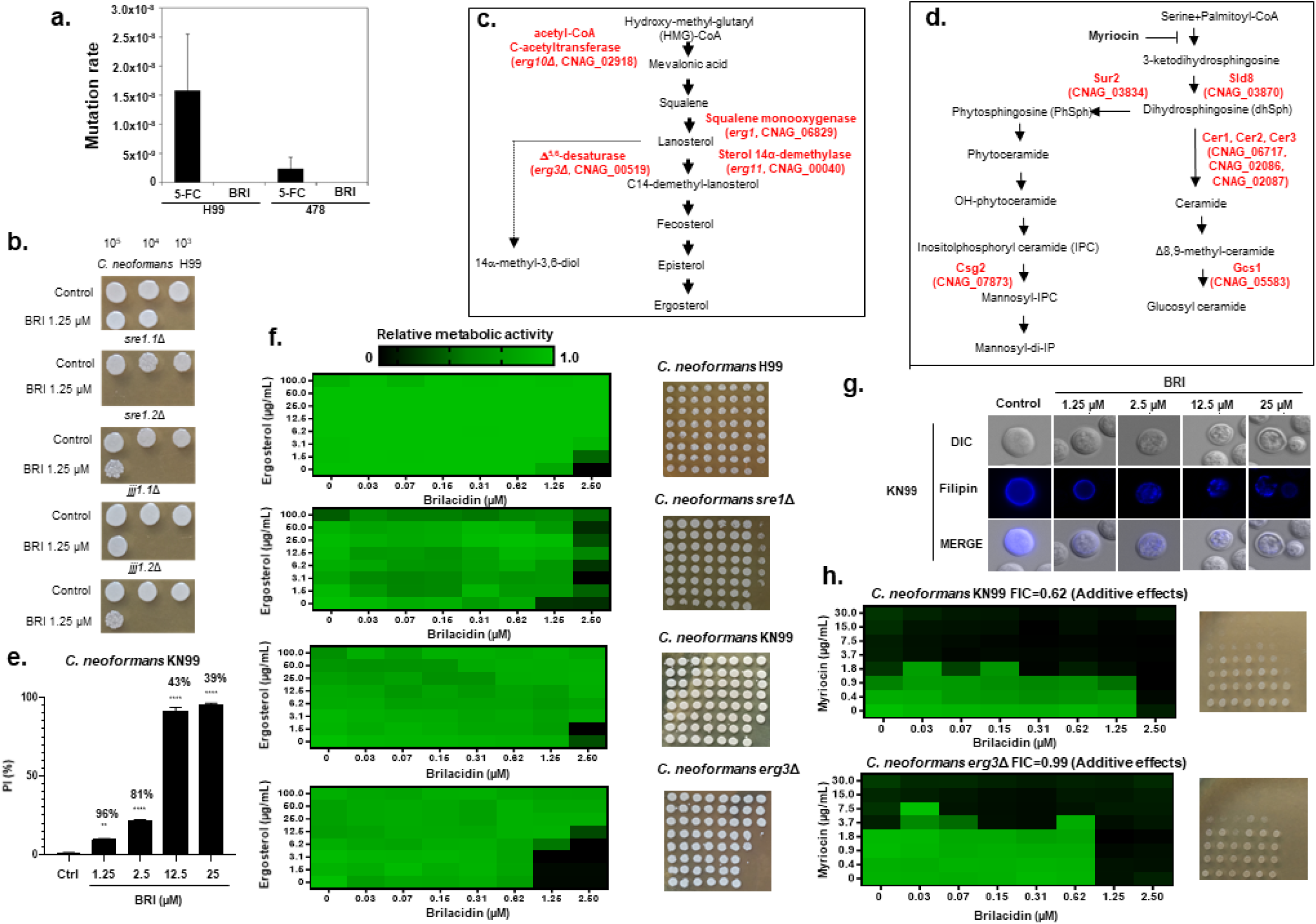
Ergosterol and glycosphingolipids are important for the BRI MoA against *C. neoformans*. **a.** Fluctuation assays for *C. neoformans* grown on solid RPMI medium containing 100 µg/mL 5-FC or 100 µM BRI. **b.** *C. neoformans* wild-type and null mutant strains were grown for 48 h in RPMI -/+ BRI (1.25 µM) and plated on solid YPD. **c. and d.** Diagrams showing the ergosterol and glycosphingolipid biosynthesis pathways in fungi. **e.** Percentage of cells with propidium iodide (PI) accumulation in the *C. neoformans* KN99a strain. Notice the % on the top of the bars represent the cell viability upon exposure to different BRI concentrations. Results shown are the average of two repetitions with 50 cells each ± standard deviation. Statistical analysis: Ordinary one-way ANOVA \*\**p value* <0.005; \*\*\*\**p value* <0.0001. **f.** XTT assays for *C. neoformans* strains grown for 48 h at 30 °C with the indicated concentrations of ergosterol and BRI. **g.** *C. neoformans* KN99a strain exposed to the indicated concentrations of BRI for 4 h and stained with filipin. **h.** Checkerboard and cidality assays for *C. neoformans* KN99a and *erg3*Δ strains grown in RPMI with the indicated concentrations of myriocin and BRI.

**Table 1.**
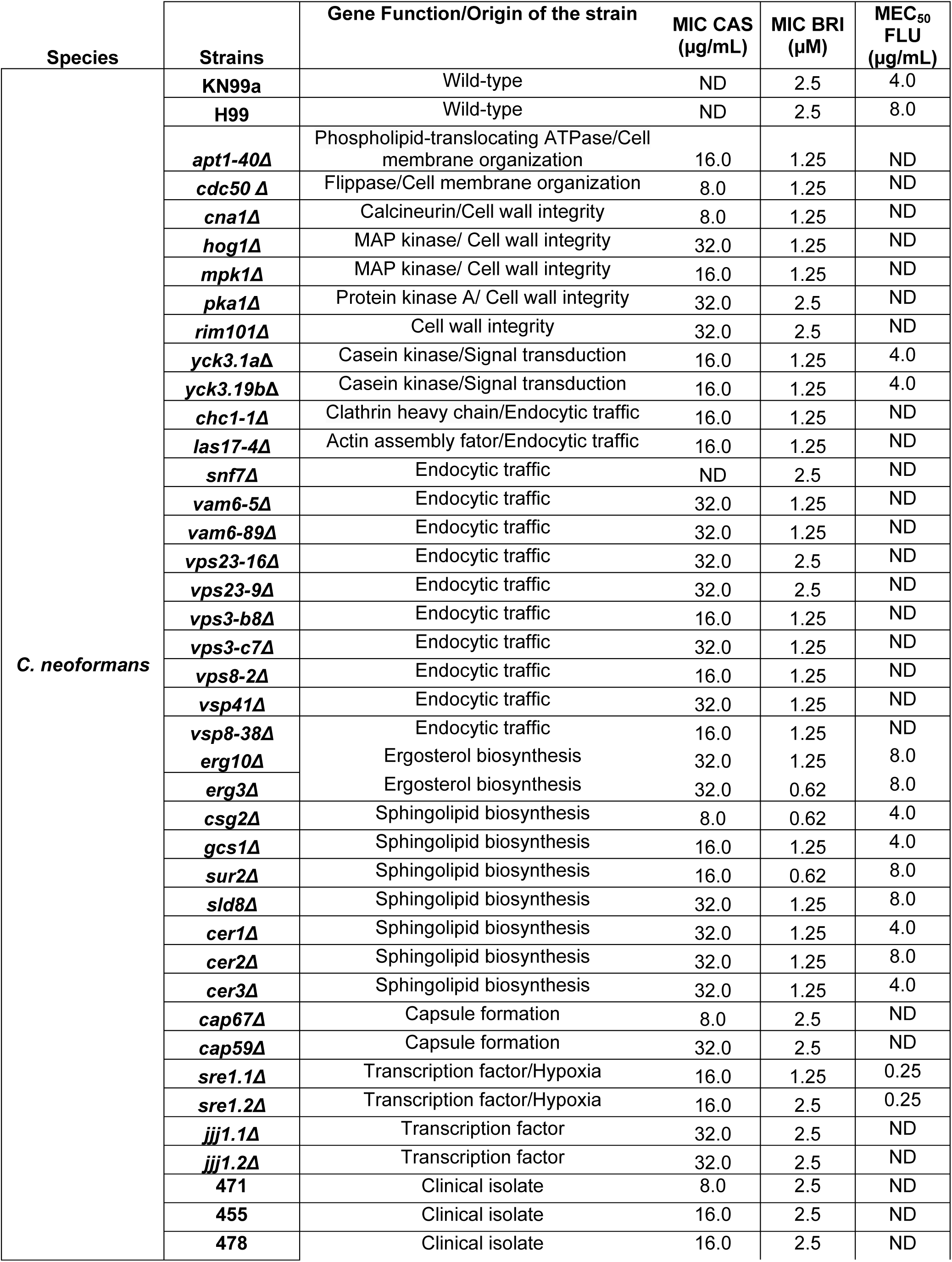
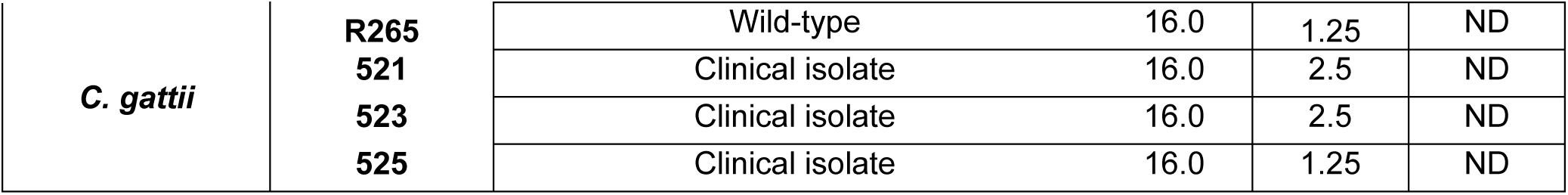
C. neoformans and *C. gattii* MICs.

As an initial step to understand BRI MoA, we screened a *C. neoformans* H99 transcription factor (TF) null mutant library^75^ for susceptibility to BRI. To do this we grew 322 signature-tagged gene deletion strains, corresponding to 155 putative TF genes, at 30 °C on liquid RPMI+BRI (1.25 µM) and then plated them on fresh solid YPD. Two TF mutants, *sre1*Δ (CNAG_04804) and *jjj1*Δ (CNAG_05538), were more susceptible to BRI than wild-type (**Figure 1b**). One of the *sre1*Δ mutants showed decreased susceptibility to BRI in solid medium but the same MIC from the wild-type strain in liquid medium (**Figure 1b and Table 1**). These two mutants were independently constructed by using biolistic bombardment and the differences between them could be due to the introduction of additional mutations in one of them. Sre1p is a homolog of the mammalian sterol regulatory element-binding protein (SREBP) that functions in oxygen-sensing^44^. *SRE1* is upregulated in the presence of fluconazole^45^, *sre1*Δ mutants are hypersensitive to azoles, and the protein is required for ergosterol biosynthesis in both hypoxic and normoxic conditions^44^. Jjj1p is a DNAJ-like co-chaperone and contains a U1 snRNA-type zinc finger; interestingly, the deletion of the *S. cerevisiae JJJ1* homolog causes defects in fluid-phase endocytosis^46^. *C. neoformans jjj1*Δ is more sensitive to AmB and more resistant to fluconazole and ketoconazole than the wild-type^47^. These results suggested a possible involvement of ergosterol in the BRI MoA.

To validate and test the involvement of *SRE1* and ergosterol, we assessed the BRI susceptibility of two null mutants in the ergosterol biosynthesis pathway: *erg3Δ* (CNAG_00519), which encodes a C-5 sterol desaturase that catalyzes the introduction of a C-5(6) double bond into episterol, a precursor in ergosterol biosynthesis, and *erg10Δ* (CNAG_02818), which encodes an acetyl-CoA C-acetyltransferase involved in the first step in mevalonate biosynthesis^48^ (**Figure 1c**). Both of them showed increaased sensitivity to BRI (**Table 1**). Loss-of-function mutations in the ergosterol biosynthetic gene *ERG3* attenuate azole toxicity in both *S. cerevisiae* and *Candida albicans*; they fail to synthesize ergosterol and exhibit increased resistance to azole antifungal drugs^49,50^. *C. neoformans erg3Δ* mutants have reduced ergosterol levels but surprisingly are more susceptible to azoles than wild-type cells^51,52^, highlighting possible metabolic differences in ergosterol biosynthesis among the three fungal systems. *C. neoformans erg3Δ* is also more sensitive to BRI than *erg10Δ*, probably related to the fact that *C. neoformans* has two *ERG10* paralogs (CNAG_02818 and CNAG_00524) (**Table 1**).

We also tested seven null mutants in the sphingolipid biosynthesis pathway^53^ (**Figure 1d**): *gcs1Δ* (CNAG_05583, glucosylceramide synthase), *csg2Δ* (CNAG_07873, inositolphosphoryl ceramide, IPC), *sur2Δ* (CNAG_03834, sphinganine C4-hydroxylase), *sld8Δ* (CNAG_03870, Δ8-fatty acid desaturase), *cer1Δ*, *cer2Δ*, and *cer3Δ* (CNAG_06717, CNAG_02086, and CNAG_02087, acyl-CoA-dependent ceramide synthases). All of these mutants showed decreased MICs for BRI (**Table 1**) when compared to the wild-type strain.

We hypothesized that the effects observed in ergosterol and sphingolipid mutants could be caused by BRI depolarizing the cell membrane, as it does in bacterial cells^54^. We attempted to determine the effect of BRI on the resting membrane potential by using the fluorescent voltage reporter DIBAC_4_(3), but this strategy did not work for *C. neoformans*. We next tested cell viability using propidium iodide (PI), a fluorescent DNA-binding dye that freely penetrates cell membranes of dead or dying cells but is excluded from viable cells. When the *C. neoformans* KN99a cells were incubated without BRI for 4 h, no cells were stained by PI, while exposure to increasing BRI concentrations (1.25 to 25 µM) yielded progressively increased staining (7 to 95%) (**Figure 1e**). This increased permeability decreased the cell viability from 96 % (1.25 µM BRI) to 39 % (25 µM BRI) (**Figure 1e**; notice the % on the top of the bars represent the cell viability upon exposure to different BRI concentrations). These results suggest that BRI induces *C. neoformans* permeabilization and cell death.

We speculated that if BRI binds directly to ergosterol, addition of ergosterol to the culture medium would prevent its activity. Ergosterol addition abrogated BRI susceptibility completely for *C. neoformans* H99 and KN99a, but to a much lesser extent for *C. neoformans sre1*Δ and *erg3*Δ (**Figure 1f**). Since these two mutants have reduced ergosterol levels, these results suggest that not all available BRI is binding to free ergosterol. Filipin is an antifungal polyene macrolide that stoichiometrically interacts with sterol to form a stable complex^55,56^. *C. neoformans* KN99a shows homogeneous filipin distribution on the cell membrane in the absence of BRI (**Figure 1g**). In contrast, increased BRI concentrations induce the formation of filipin stained patches on the cell membrane (**Figure 1g**), suggesting BRI affects the ergosterol distribution and organization on the cell membrane.

To determine whether BRI displayed any interaction with sphingolipid, we combined various concentrations of BRI and myriocin, an inhibitor of serine palmitoyltransferase (**Figure 1d**), the first step in glycosphingosine biosynthesis^57^. Checkerboard assays showed FICs of 0.62 and 0.99 for both wild-type and *erg3*Δ strains, respectively, indicating an additive effect between BRI and myriocin (**Figure 1h**).

We also examined the expression of genes involved in ergosterol biosynthesis (**Figure 1c**) when *C. neoformans* was grown for 4 h in the presence of BRI (25 µM) or BRI+CAS (25 µM+8 µg/mL). Compared to growth with no drug or CAS alone, *ERG3* and *SRE1* expression was induced by BRI about 10- to 15-fold, respectively, while *ERG1* and *ERG11* expression did not change (**Figures 1c and 2a**). However, these levels of exposure to BRI did not modify the ergosterol content of the H99 wild-type and *erg3*Δ strains (notice that *erg3*Δ has lower ergosterol than the wild-type in the absence of BRI; **Figure 1c and 2b**).

**Figure 2.**
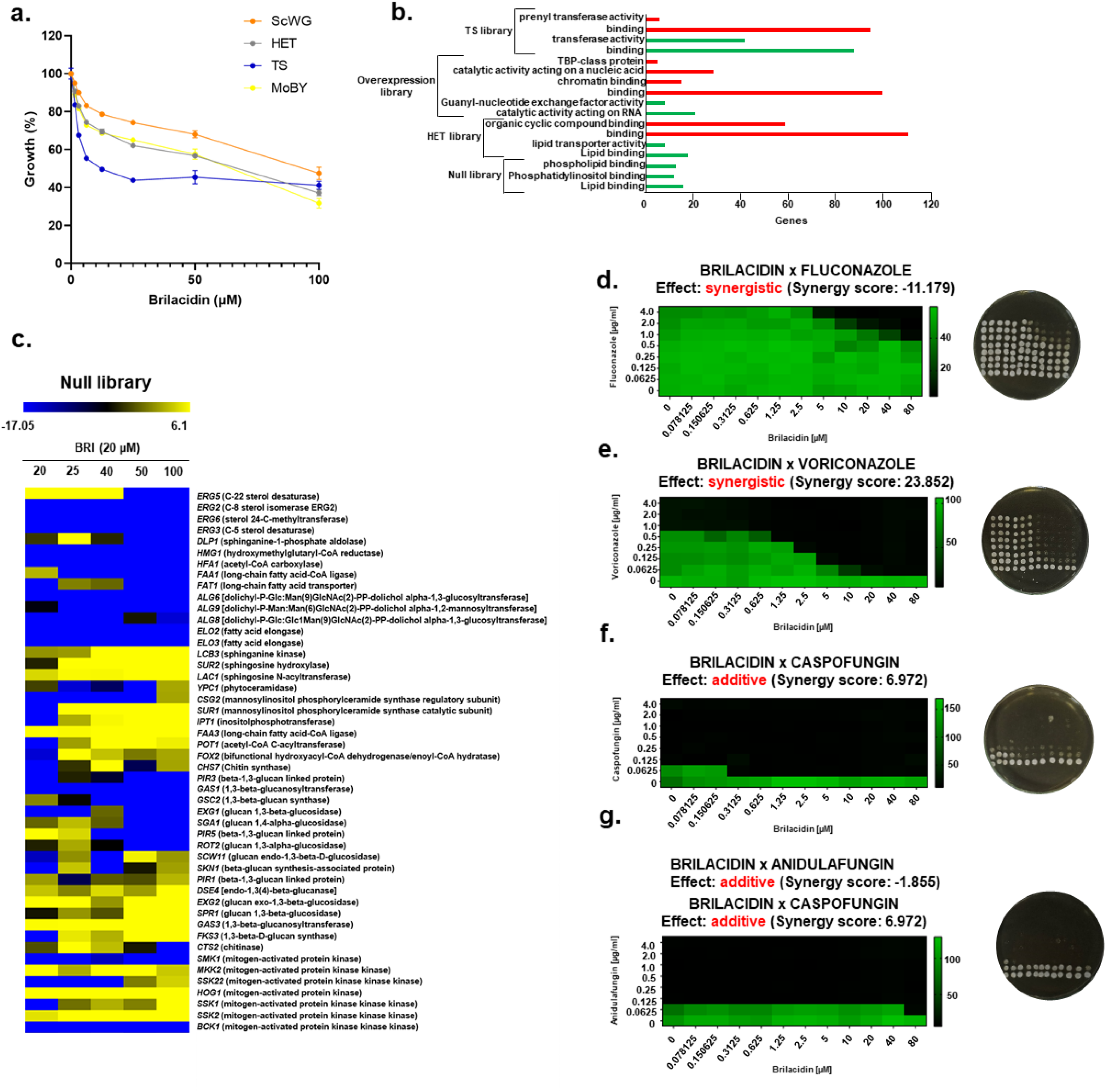
*C. neoformans* retrograde sterol transporter *ysp2*Δ leads to abnormal accumulation of ergosterol at the plasma membrane and it is less susceptible to CAS and BRI. **a.** RTqPCR for *ERG1*, *ERG3*, *ERG11*, and *SRE1* mRNA accumulation when *C. neoformans* H99a wild-type was grown for 16 h and transferred or not to either BRI (25 µM), CAS (8 µg/mL), or BRI+CAS (25 µM+8 µg/mL). The results are the average of three repetitions ± standard deviation. Statistical analysis: 2way ANOVA \*\*\*\**p value* < 0.0001. Cells lacking Ysp2 exhibit enhanced resiliance to CAS and BRI. B. Ergosterol concentration in the *C. neoformans* H99a wild-type and *erg3*Δ mutant. **c. and d.** Wild-type strain (KN99a) or *ysp2*Δ cells were grown in YNB (pH 7.4) for 72 h at 37°C with CAS (**c.**) or BRI. (**d.**) and plated for viability. Mean ± SE values of colony forming units (CFUs) per well are plotted; shown is one representative study of three independent experiments performed in technical triplicate. Statistical analysis: unpaired Student’s t-test *p < 0.05; **p< 0.005. **e. and f.** The same strains were grown in YNB (pH 7.4) for 48 h at 30°C in the presence of the indicated concentrations of CAS and BRI. Metabolic activity measured by XTT assay is shown. ΣFIC, combined fractional inhibitory concentration; black boxes, Abs_492_ > 1.0.

The *YSP*2 gene of *C. neoformans* encodes a retrograde sterol transporter^52^. Cells lacking this gene (*ysp2*Δ) accumulate ergosterol at the plasma membrane, leading to dramatic deformations of the membrane and cell wall. These mutant cells are also more sensitive than wild-type to the sterol-targeting antifungal compound AmB and other membrane stressors^52^. To measure the response of this mutant to BRI and CAS, we performed MIC assays, using YNB medium because *ysp2*Δ cells grow extremely poorly in RPMI^52^. We found that the MIC for each compound was higher for *ysp2*Δ than for the corresponding wild-type, KN99a: 32 versus 8 µg/mL for CAS (**Figure 2c**) and 2.5 versus 1.25 µM for BRI (**Figure 2d**). We noted that the MIC for CAS and BRI in KN99a strain grown in RPMI was higher, >32 µg/mL and 2.5 µM, respectively, than in YNB (see Table 1). Next, we tested for potential interactions of these compounds by performing checkerboard assays, again using YNB medium. These experiments indicated an additive effect, with no synergy or antagonism between the two drugs for either KN99a or *ysp2*Δ cells (**Figures 2e and 2f**).

Taken together, these results suggest that the primary BRI MoA in *C. neoformans* is perturbation of the cell membrane that reduces its organization and integrity, leading to increased membrane permeability and cell death. Although our results suggest that BRI can bind to free ergosterol, reduced sphingolipid concentration in the cell membrane also potentiates the MoA of BRI against *C. neoformans*.

### BRI affects cell membrane organization, the CWI pathway and calcium metabolism in *S. cerevisiae*

As an additional tool to investigate BRI MoA, we performed several screens using haploid null (ScWG), temperature-sensitive (TS), overexpression (MoBY), and heterozygous (haploinsufficiency profiling, HET) strain libraries of the model yeast *S. cerevisiae*^22,58^. Pooled cultures of barcoded *S. cerevisiae* strains from these collections were grown in liquid YPGal medium for 19 to 28 h at 26 or 30 °C (**Figure 3a**). The relative abundance of each barcoded strain following BRI treatment was then determined by highthroughput sequencing of PCR-amplified barcodes followed by BEAN-counter analysis^59^ to measure enrichment or depletion of each strain in the presence of BRI relative to solvent control (**Supplementary Tables S1 to S4 and Figures S1 to S4; 10.6084/m9.figshare.25239550**). For all these analyses, we considered as enriched or depleted log2 values of ≥ 2 or ≤ −2, respectively.

**Figure 3.**
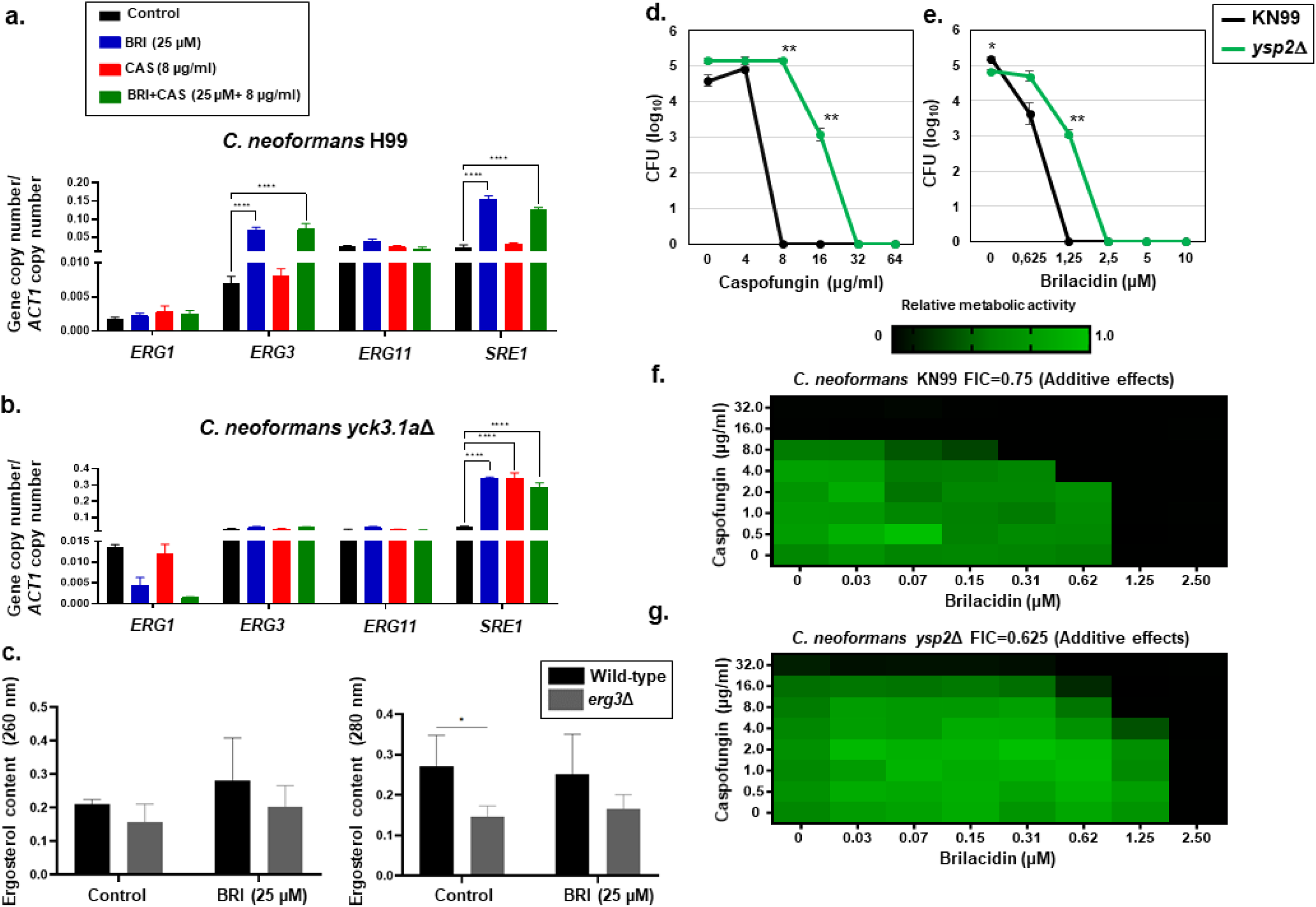
In *S. cerevisiae*, BRI affects cell membrane organization, cell wall integrity pathway and calcium metabolism. **a.** *S. cerevisiae* pooled haploid null (ScWG), heterozygous (HET), temperature sensitive (TS), and overexpression (MoBY) libraries were grown in liquid YPGal medium in the absence or presence of BRI for 19 h at 30 °C (ScWG and HET, 6.25 to 100 µM BRI), 28 h at 26 °C (TS, 1.5625 to 25 µM BRI), or 24 h at 30 °C (MoBY, 20 to 100 µM BRI). **b.** Categorization of the hits for each library according to Gene Ontology Term Finder (https://www.yeastgenome.org/goTermFinder) using a *p-* corrected value < 0.001. The number of engineered genes is indicated for each library as red for depleted or green for enriched. **c.** Heat map of the genes that correspond to the mutants depleted or enriched in the presence of BRI and visually inspected as more modulated. **d. to g.** Checkerboard and cidality assays for *S. cerevisiae* grown in YPGal with the indicated concentrations of BRI+Fluconazole, BRI+Voriconazole, BRI+CAS, and BRI+Anidulafungin.

Of 3,642 genes from the *S. cerevisiae* haploid null library, the normalized log read count profile showed 323 mutants that were depleted and 426 that were enriched when grown in the presence of several concentrations of BRI, more specifically 100 µM (**Supplementary Table S1 and Figure S1; 10.6084/m9.figshare.25239550**). Among the top 200 mutants depleted, we observed Gene Ontology (GO) enrichment for lipid binding, phosphatidylinositol binding, and phospholipid binding (**Figure 3b**). There was no category enrichment for the enriched mutants, but the 10 most enriched included *SNF1* and *SNF4* (encoding AMP-activated serine/threonine-protein kinase catalytic and regulatory subunits, respectively, required for glucose-shortage induced respiratory gene expression), *SSK2* (a MAP kinase kinase kinase of the *HOG1* mitogen-activated signaling pathway), and *SUR1*, *SUR2*, and *IPT1* (encoding a mannosylinositol phosphorylceramide synthase catalytic subunit, a sphinganine C4-hydroxylase, and inositolphosphotransferase, all required for sphingolipid biosyntheis) (**Figure 3b and Supplementary Table S1 and Figure S1; 10.6084/m9.figshare.25239550**). We also observed *ERG3*, *ERG5*, *ERG2*, and *ERG6* (encoding C-5 and C-22 sterol desaturases, C-8 sterol isomerase, and sterol 24-C-methyltransferase) mutants as depleted (**Supplementary Table S1 and Figure S1; 10.6084/m9.figshare.25239550**), supporting the involvement of ergosterol biosynthesis in the BRI MoA. The mitogen-activated protein (MAP) kinase Kinase Kinase (KKK) BCK1 and MAPKK MKK1 mutants were depleted while the MAPKK MKK2 mutant was enriched, suggesting the involvement of the protein kinase C signaling and CWI in the BRI MoA (**Supplementary Table S1 and Figure S1; 10.6084/m9.figshare.25239550**).

In the HET analysis, there were 68 mutants depleted with GO enrichment for protein binding and 99 mutants enriched which showed GO enrichment for catalytic binding on nucleic acid and RNA (**Figure 3b, Supplementary Table S2 and Figure S2; 10.6084/m9.figshare.25239550**). Interestingly, a mutant in *CMD1* (encoding calmodulin) was the third most depleted mutant in the HET (**Supplementary Table S2 and Figure S2; 10.6084/m9.figshare.25239550**).

In the overexpression library, we identified 59 strains that were depleted. These were enriched for GO categories such as catalytic activity acting on RNA and guanyl-nucleotide exchange factor activity. Fifty-six others showed enhanced growth in the presence of BRI, but with no significant category enrichment (**Figure 3b**, **Supplementary Table S3 and Figure S3; 10.6084/m9.figshare.25239550**). Interestingly, the overpression of *ERG25* and *ERG20* [(encoding a methylsterol monooxygenase and a bifunctional (2E,6E)-farnesyl diphosphate synthase, respectively)] increased survival of these strains in the presence of BRI, once more suggesting that the modulation of the ergosterol pathway is important for BRI MoA (**Figure 3b**, **Supplementary Table S3 and Figure S3; 10.6084/m9.figshare.25239550**). In the TS library, there were 141 depleted mutants enriched for GO category of protein binding and 76 enriched mutants with GO category transferase activity (**Figure 3b**, **Supplementary Table S4 and Figure S4; 10.6084/m9.figshare.25239550**). *CMD1* was the top depleted mutant in this TS list, again highlighting the important role of calcium metabolism in BRI MoA (**Figure 3b**, **Supplementary Table S4 and Figure S4; 10.6084/m9.figshare.25239550**). Surprisingly, the overexpression of *PKC1* (encoding a protein serine/threonine kinase that is essential for cell wall remodeling during growth) increases the mutant depletion, again implicating the CWI pathway in the MoA of BRI against *S. cerevisiae* (**Supplementary Table S3 and Figure S3; 10.6084/m9.figshare.25239550**).

We also examined combinations of antifungal compounds. Since the BRI was not able to inhibit *S. cereviase* growth at the highest concentration tested, the FIC value was not calculated. On the other hand, we used the SynergyFinder2.0 (https://synergyfinder.fimm.fi) to calculate the interaction score between BRI and the other antifungal compounds. The interaction between the drugs was classified based on the synergy score where values lower than −10 were considered antagonistic, values from −10 to 10 was considered additive and values higher than 10 were considered synergistic. BRI potentiated azoles with cidal effects against *S. cerevisiae*, with scores of 23.854 and 23.852 for fluconazole and voriconazole, respectively, and additive synergy scores of 6.972 and −1.855 with CAS and anidulafungin, respectively (**Figure 3c to 3f**).

Taken together, these results suggest that the MoA of BRI in *S. cerevisiae* involves ergosterol and sphingolipids biosynthesis, the CWI pathway, and calcium metabolism.

### BRI potentiates AmB against *C. neoformans*

We have previously demonstrated that BRI potentiates CAS activity against *C. neoformans*^41^. Checkerboard assays showed that BRI+CAS had an additive interaction against *C. neoformans* strains H99 and KN99a and *C. gattii* strain R265 (FICs of 0.56, 0.56, and 0.76, respectively; **Figures 4a, 4b, and 4c**). In contrast, we did not observe any interaction between BRI+micafungin and BRI+anidulafungin (**Figures 4d and 4e**). There is an additive interaction between BRI+AmB, BRI+voriconazole, BRI+fluconazole, and BRI+5-FC (**Figure 4d to 4i**) against *C. neoformans*. CAS affects β-1,3-glucan biosynthesis activating cell wall assembly and remodeling, preventing resistance against osmotic forces, which then leads to cell lysis. However, addition of different concentrations of sorbitol cannot improve *C. neoformans* growth in the presence of BRI (**Supplementary Figure S5**, **10.6084/m9.figshare.25239550;** BRI and different concentrations of sorbitol show a FIC index >1, indifference).

**Figure 4.**
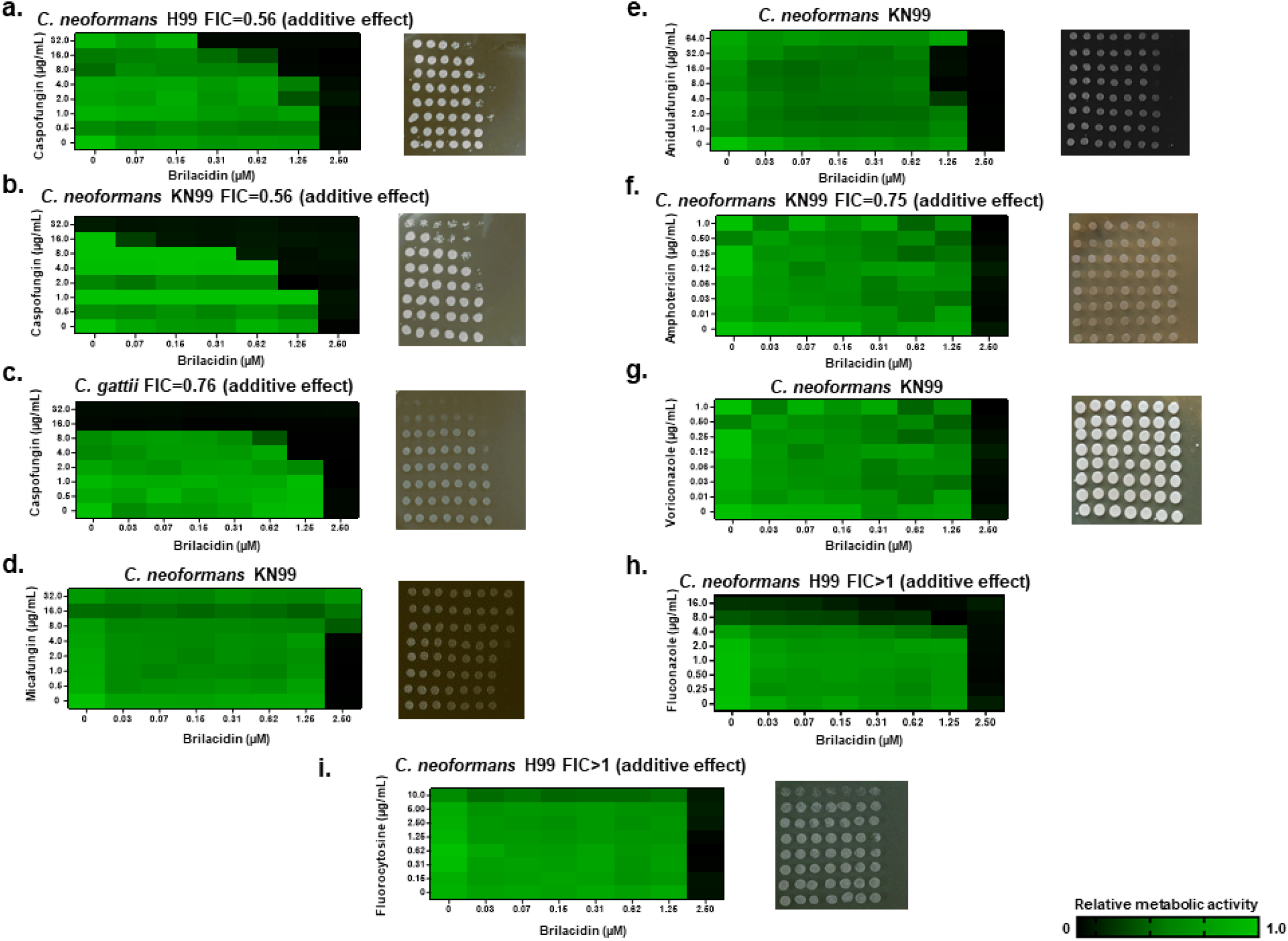
BRI potentiates CAS against *C. neoformans* and *C. gattii.* **a to c.** Checkerboard and cidality assays using XTT. *C. neoformans* and *C. gattii* were grown for 48 h in RPMI medium with various combinations of BRI+CAS. **d.** BRI+Micafungin. **e.** BRI+Anidulafungin. **f.** BRI+AmB. **g.** BRI+Voriconazole. **h.** BRI+Fluconazole. **i.** BRI+5_FC.

### Casein kinases are important for BRI and CAS activities

To further assess the BRI MoA, a collection of 58 protein kinase inhibitors (PKI, at a concentration of 20 µM; **Supplementary Table S5; 10.6084/m9.figshare.25239550**) was screened for effects on *C. neoformans* H99 growth and corresponding metabolic activity alone or together with BRI (1.25 µM) (**Figure 5a and Supplementary Table S5; 10.6084/m9.figshare.25239550**). Six PKIs [compounds 3, 19, 27, 41, 53, and 58, which correspond to FRAX486 (targets p21-activated kinase, PAK), PP121 (targets STK25), AT7867 (an ATP-competitive inhibitor of Akt1/2/3 and p70S6K/PKA), refametinib (targets MAP2K2), GW461487A (targets p38 kinase), and GW431756X (whose target is unknown), respectively] completely inhibited *C. neoformans* growth. Three casein kinase inhibitors, PF-670462 (compound 34), UNC-ALM-39 (compound 55), and UNC-ALM-87 (compound 57), all (https://www.selleckchem.com/products/pf-670462.html;^60^), potentiated BRI activity against *C. neoformans*^60^ (**Figures 5a to 5c**).

**Figure 5.**
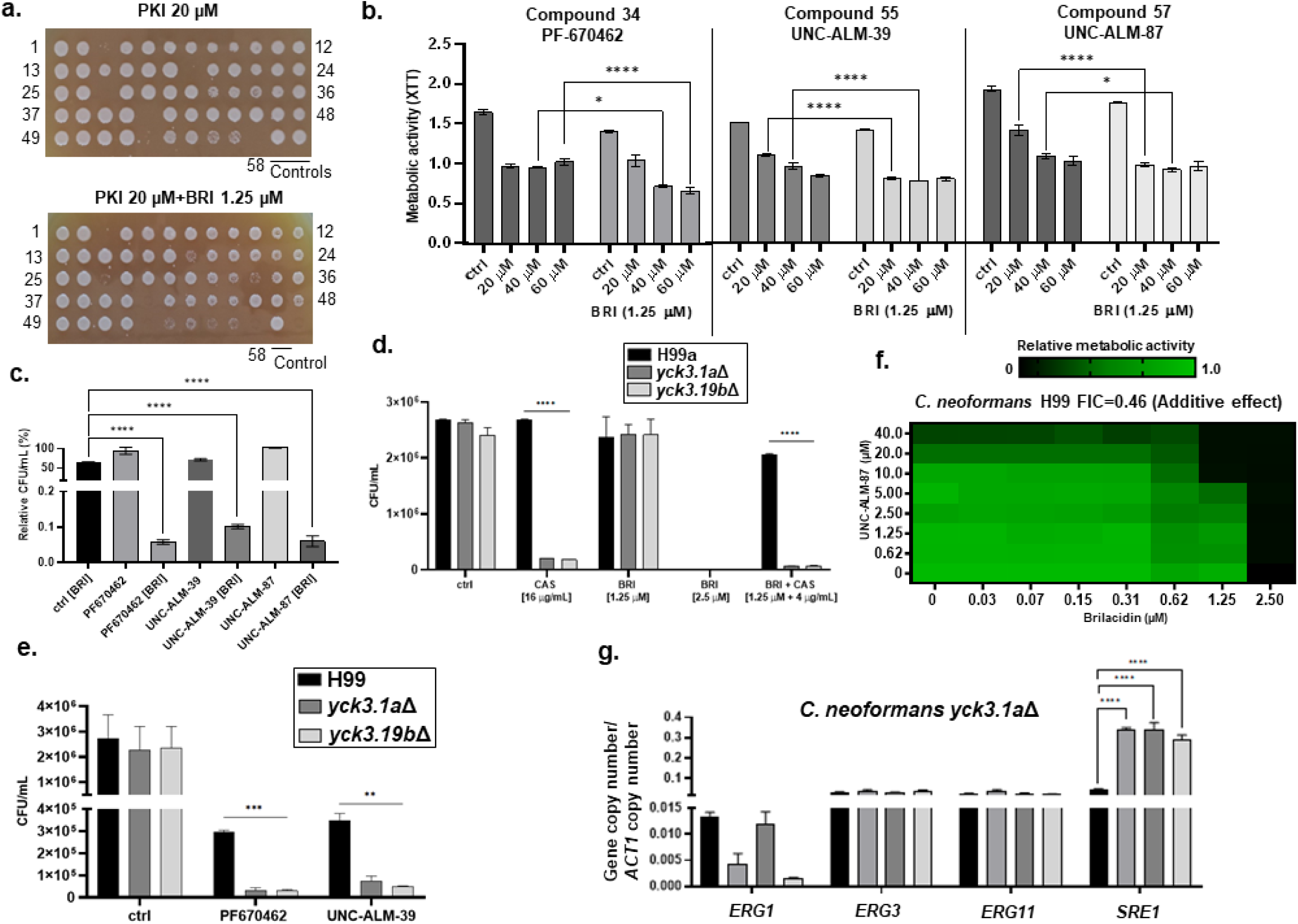
Casein kinases are involved in BRI MoA. **a.** *C. neoformans* growth screening of 58 PKIs (20 µM) in the absence or presence of BRI (1.25 µM). Cells were grown for 48 h at 30 °C in RPMI medium and plated on solid YPD medium. **b.** Reduction of metabolic activity by PKIs in the absence or presence of BRI. The cells were grown for 48 h at 30 °C in RPMI medium and metabolic activity was measured by using XTT. **c.** Viability of *C. neoformans* growth with or without PKIs for 48 h at 30 °C. **d.** Viability of *C. neoformans* wild-type and *ypk3*Δ mutants after incubation for 48 h at 30 °C in RPMI medium in the absence or presence of various concentrations of BRI and/or CAS. **e.** Viability of *C. neoformans* wild-type and *yck3*Δ mutants after incubation for 48 h at 30 °C in RPMI medium in the absence or presence of PKIs. Statistical analysis: 2way ANOVA \**p value* <0.05; \*\**p value <0.01;* \*\*\**p value* <0.001; \*\*\*\**p value* <0.0001 **f.** Checkerboard assay using XTT. *C. neoformans* was grown for 48 h in RPMI medium with various combinations of UNC-ALM-87 and BRI.

*C. neoformans* encodes two casein kinase 1 (CK1) paralogs, *CCK1* and *CCK2*^61^. *CCK2* is an essential gene while *CCK1* mutants are hypersensitive to sodium dodecyl sulfate (SDS) treatment, osmotic and oxidative stresses, AmB, fluconazole, Congo red, calcofluor white, and regulates the phosphorylation of both Mpk1 and Hog1 mitogen-activated protein kinases during these stresses^61,62^. Two independent *cck1* (here called *yck3Δ*) mutants in the *C. neoformans* H99 background were more sensitive than wild-type to CAS and BRI+CAS but surprisingly not to BRI alone (**Figure 5d**). The differences between the casein kinase PKIs potentiating BRI against *C. neoformans* and the lack of increased susceptibility of the *yck3Δ* mutants to BRI could be explained if (i) the three PKIs do not inhibit *C. neoformans* Yck3 or (ii) Cck2p is more important than Yck3 for the BRI mechanism of action. To address the first hypothesis, we incubated wild-type and *yck3Δ* mutant strains with the PKIs and assessed their viability (**Figure 5e**). The *yck3Δ* mutants were more sensitive to the PKIs than the corresponding wild-type strain (**Figure 5e**), emphasizing that *C. neoformans* casein kinases are targets for these compounds. Checkerboard assays confirmed an additive interaction between one of these PKIs (UNC-ALM-87) and BRI (**Figure 5f**).

We next examined the expression of genes involved in ergosterol synthesis in *C. neoformans yck3*Δ grown for 4 h in the presence of BRI and/or CAS. BRI (25 µM) and BRI+CAS (25 µM+8 µg/mL) for 4 h induce about 7-fold more *ERG1* than the wild-type strain in the control and in the presence of CAS with 3-fold reduction, but still higher levels than the wild-type, in the presence of BRI (**Figures 2a and 5g**). *ERG3* and *ERG11* showed no induction in the presence of the drugs but in the control both gene levels are about 50% higher than in the wild-type strain (**Figures 2a and 5g**). *SRE1* expression is about 50% higher in the control, BRI and BRI+CAS than the wild-type strain, and additionally it was induced in the presence of CAS alone, which was not observed in the wild-type strain (**Figures 2a and 5g**).

Taken together, these results strongly suggest that casein kinases are involved in the BRI MoA and are important for the modulation of the expression of genes involved in the ergosterol biosynthesis.

### BRI potentiates CAS against *C. neoformans* through multiple mechanisms

To explore how BRI impacts the effect of CAS on *C. neoformans*, we examined the β −1,3-glucan synthase (Fks1) that is targeted by CAS. We observed without any further quantification that cell surface levels of Fks1:GFP were apparently lower in the presence of CAS-F (a functional fluorescently labeled probe;^63^) and BRI+CAS-F than in the control (**Figure 6a**). We hypothesized that this reflected different levels of endocytic removal of the Fks1:GFP. When *C. neoformans* is treated either with CAS-F or BRI+CAS-F, both Fk1:GFP and CAS-F are internalized into small vesicles and structures that resemble vacuoles (**Figure 6a**). The vacuoles were confirmed by co-staining with CellTracker Blue^TM^ CMAC Dye (7-amino-4-chloromethylcoumarin), a hydrophobic molecule that enters the fungal cytoplasm and is modified through binding to glutathione by glutathione S-transferase^64^. In fungi, CMAC accumulates in vacuoles, most likely through the transport of glutathione transporters present on the vacuolar membrane^64^. Unfortunately, in spite of several attempts, the resolution of the microscopy did not allow us to perform a differencial analysis of the number of vesicles/vacuoles populated by CAS-F upon BRI treatment. However, our results suggest a possible internalization of Fks1:GFP and CAS-Fks1:GFP complexes by the *C. neoformans* endocytic machinery, followed by trafficking to the vacuoles.

**Figure 6.**
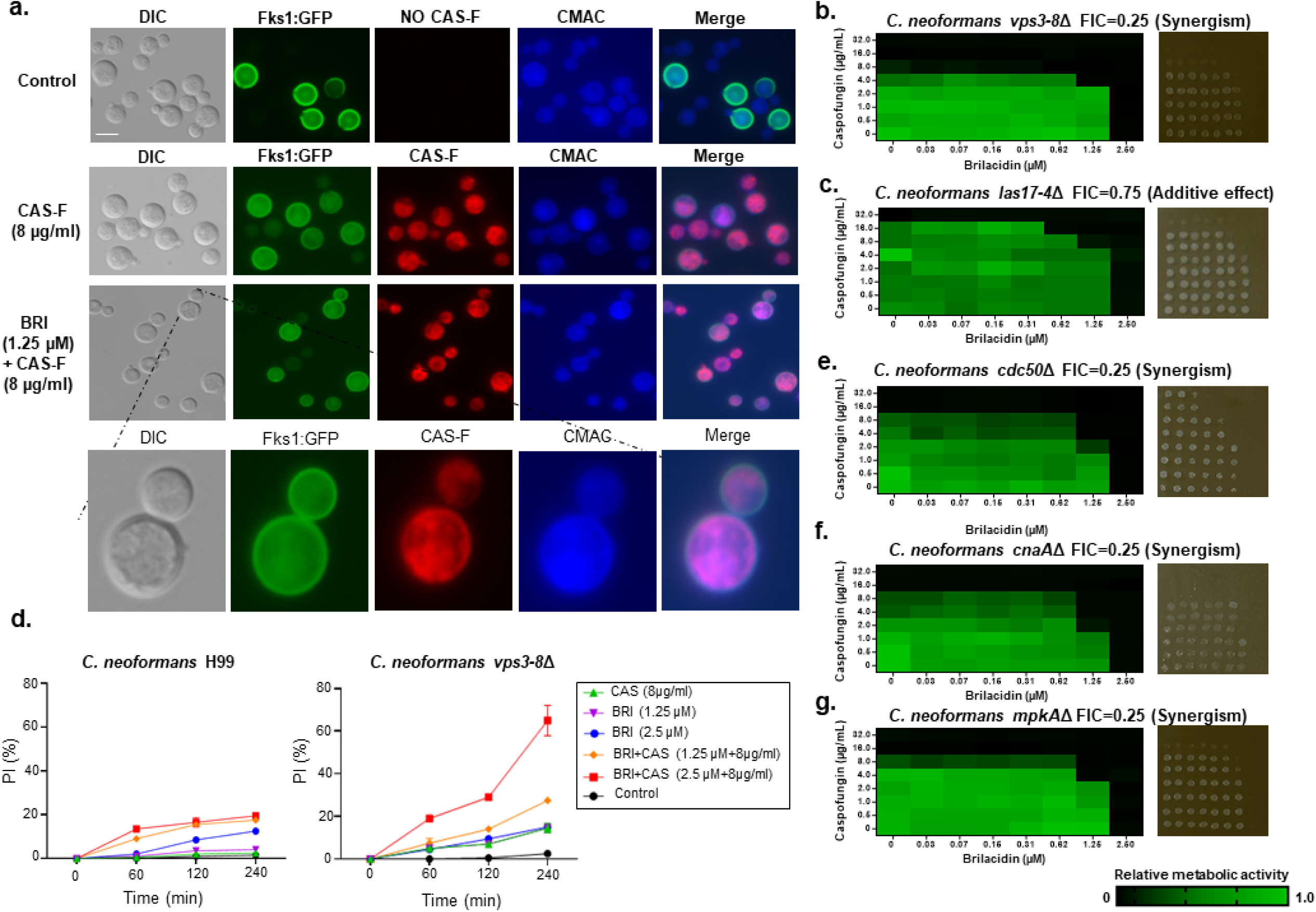
Potentiation of CAS by BRI against *C. neoformans* depends on the endocytic pathway. **(a)** *C. neoformans* Fks1:GFP was grown for 16 h in YPD at 30 °C and incubated in RPMI medium with or without CAS-F (fluorescent caspofungin) or BRI+CAS-F. CMAC, CellTracker Blue CMAC Dye (7-amino-4-chloromethylcoumarin. **b and c.** Checkerboard and cidality assays using XTT. *C. neoformans vps3-8*Δ **(b)** and *las17-4*Δ **(c)** were grown for 48 h at 30 °C with various concentrations of BRI and CAS. **(d)** Percentage of cells with propidium iodide (PI) accumulation in *C. neoformans* H99 and *vps3-8*Δ strains. The cells were exposed to different concentrations of BRI for 4 h and cells with PI accumulated in the cytoplasm were counted. The results are the average of two repetitions with 50 cells each ± standard deviation. **(e to g)** Checkerboard and cidality assays using XTT. *C. neoformans cdc50*Δ **(e)**, *cna1*Δ **(f)**, and *mpk1*Δ **(g)** were grown for 48 h at 30 °C with different concentrations of BRI and CAS.

To test this hypothesis, we selected a group of mutants lacking genes involved in the endocytic machinery (see **Table 1** and ^65–67^; for reviews ^68–70^) and measured their MICs in the presence of BRI or CAS (**Table 1**). For these studies we chose (i) two members of the CORVET membrane tethering complex (two independent null mutants of *VPS3*, encoding a cytoplasmic protein required for the sorting and processing of soluble vacuolar proteins, *vps3-c7*Δ and *vps3-b8*Δ and two independent null mutants of *VPS8*, encoding a membrane-binding component of the CORVET complex, *vps8-2*Δ and *vps8-38*Δ), (ii) one member of the ESCRT-I complex (two independent null mutants of *VPS23*, involved in ubiquitin-dependent sorting of proteins into the endosome, *vps23-9*Δ and *vps23-16*Δ), (iii) one member of the ESCRT-III complex (*SNF7*, involved in the sorting of transmembrane proteins into the multivesicular body (MVB) pathway), (iv) two members of the HOPS endocytic tethering complex (two independent null mutants of *VAM6*, encoding a guanine nucleotide exchange factor for the GTPase Gtr1p, *vam6-5*Δ and *vam6-89*Δ) and *VPS41* (encoding a vacuole membrane protein that functions as a Rab GTPase effector), (v) *CHC1* (encoding a clathrin heavy chain, *chc1-1Δ*), and (vi) *LAS17* (encoding an actin assembly factor, *las17-4*Δ). Apart from *vps23*Δ and *snf7*Δ, all mutants showed reduced MICs for BRI and CAS (**Table 1**). Checkerboard assays showed that BRI and CAS have synergism and an additive interaction against *C. neoformans vps3-8*Δ and *las17-4*Δ mutant strains with FICs of 0.25 and 0.75, respectively (**Figures 6b and 6c**). PI permeability into the cytoplasm is increased in the *vps3-8*Δ mutant when compared to the wild-type strains: BRI 1.25 and 2.5 µM, 10 % versus 2 and 10 %; CAS 8 µg/mL, 0 % versus 2 %; BRI+CAS (1.25 µM+8µg/mL), 25 % versus 16 %; and BRI+CAS (2.5 µM+8µg/mL), 65 % versus 17 % (**Figure 6d**). Recently, *C. neoformans CDC50*, encoding a lipid flippase responsible for the translocation of phospholipids from the exocytoplasmic to the cytoplasmic leaflet, was shown to be important for CAS resistance. Loss of this leads to abnormal phospholipid distribution and impaired intracellular vesicular trafficking^37,71^. We observed that *cdc50*Δ is not only more sensitive than wild type to CAS but also to BRI (**Table 1**). Checkerboard experiments with *cdc50*Δ showed synergism between CAS and BRI (FIC of 0.25; **Figure 6e**).

Previously, we have shown that BRI acts in *A. fumigatus* in part by affecting CWI pathway^41^. We therefore tested the BRI and CAS susceptibility of a group of *C. neoformans* null mutants involved in the CWI pathway: (i) *CNA1* (encoding a catalytic subunit of the phosphatase calcineurin), (ii) *HOG1* and *MPK1* (encoding mitogen-activated protein kinases from the high-osmolarity glycerol pathway and CWI, respectively), (iii) *PKA1* (encoding the catalytic subunit of the protein kinase A), and (iv) *RIM101* (encoding a transcription factor responsible for sensing extracellular pH signals) (for reviews, see ^72,73^). All of these mutants showed decreased CAS MICs, consistent with prior reports on *cna1*Δ and *mpk1*Δ^32,74^. All except *rim101*Δ showed decreased MICs for BRI (**Table 1**). Checkerboard assays showed that BRI and CAS are synergistic against *C. neoformans cna1*Δ and *mpk1*Δ with FICs of 0.25 for both mutant strains (**Figures 6b and 6c**).

The cryptococcal cell wall is arranged in two layers that show different electron densities by transmission electron microscopy (TEM) (**Figure 7a**). The inner layer is composed of β-glucans and chitin, mannoproteins and melanin, while the outer layer mainly contains α- and β-glucans (Agustinho *et al.*, 2018). Treatment with CAS (8 µg/mL) led to increases in both the total cell wall and the inner layer, while BRI (25 µM) alone or BRI+CAS (25 µM + 8 µg/mL) yielded overall thickness similar to wild type but marked reduction in the inner layer compared to both wild type and CAS treatment alone (**Figures 7a, 7b and 7c**). To pursue these results, we examined the expression of genes involved in wall synthesis. We found that expression of *FKS1* was induced about 10-, 4-, and 5-fold, respectively, when *C. neoformans* was grown in the presence of BRI, CAS, and BRI+CAS for 4 h (**Figure 7d**). Expression of *CHS3* (encoding a chitin synthase) was induced about 2.5-fold when *C. neoformans* was exposed to either BRI or BRI+CAS while expression of *CHS4* and *CHS6* (encoding chitin synthases) rose 100-, 2-, and 40-fold 2.5-, 7-, and 2.5-fold, respectively, in the presence of BRI, CAS, or CAS+BRI (**Figure 7d**). Similar analysis of *C. neoformans yck3*Δ showed no induction of *FKS1* with BRI, CAS, or BRI+CAS for 4 h although *FKS1* expression was about 10-fold higher in the control than in the wild-type (**Figure 7e**); *CHS3* showed about 3-fold induction in the presence of CAS but not BRI and BRI+CAS (**Figure 7e**). *CHS4* was induced in all treatments but about 7-fold more in the control than in the wild-type control, while *CHS6* had about 4-fold induction in the control than in the wild-type control, but induction only in the presence of CAS (**Figure 7e**).

**Figure 7.**
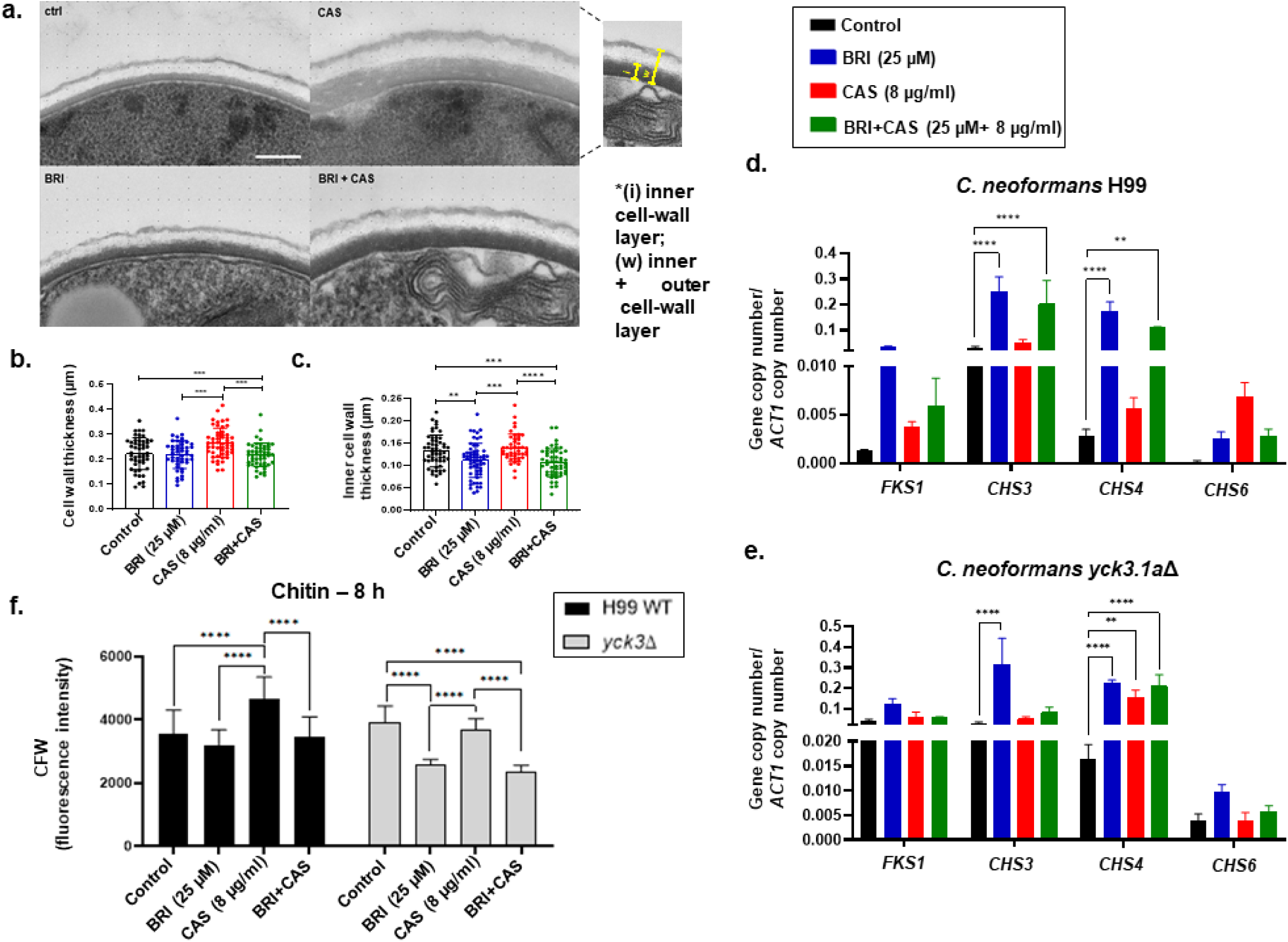
The *C. neoformans* cell wall integrity (CWI) pathway is important for the BRI+CAS MoA. **a.** Representative transmission electron microscopy (TEM) images of *C. neoformans* cells grown with no drug, BRI (25 µM), CAS (8 µg/mL), or BRI+CAS (25 µM+8 µg/mL) for 8 h. All images are to the same scale and the magnification is 3000 x; scale bar, 2 µm. Thickness of the full cell wall **(b)** and inner cell wall layer **(c)** of *C. neoformans* cells exposed to no drug, BRI (25 µM), CAS (8 µg/mL), or BRI+CAS (25 µM+8 µg/mL) for 8 h. The regions measured are indicated at the right of panel **a** and the results shown are derived from 50 *C. neoformans* cells for each treatment. Statistical analysis: Ordinary one-way ANOVA \**p value* <0.05; \*\*\**p value* <0.001; \*\*\*\**p value* <0.0001. **d. and e.** RTqPCR for *FKS1*, *CHS3*, *CHS4*, and *CHS6* mRNA accumulation when *C. neoformans* H99 wild-type and *yck3*Δ were grown for 16 h and transferred or not to either BRI (25 µM), CAS (8 µg/mL), or BRI+CAS (25 µM+8 µg/mL). The results are the average of three repetitions ± standard deviation. Statistical analysis: 2way ANOVA \*\**p value* < 0.01, and \*\*\*\**p value* < 0.0001.

Next, we evaluated whether BRI or BRI+CAS could impact chitin distribution, organization, and exposure in the cell wall by staining it with the fluorescent dye Calcofluor White (CFW) (**Figure 7f**). Exposure of *C. neoformans* wild-type or *yck3*Δ cells to CAS for 8 h increased chitin staining on the cell surface, but this effect can be decreased by BRI and BRI+CAS (**Figure 7f**). Interestingly, although chitin synthase levels are higher in the *yck3*Δ mutant than in the wild-type, the chitin levels are lower in the mutant than in the wild-type in all the treatments, except in the control, where the chitin levels are higher than in wild-type (**Figures 7d, 7e, and 7f**).

Taken together, these results suggest that the endocytic machinery plays a role in the MoA of BRI+CAS combinations, affecting cell permeability. Moreover, the CWI pathway is also important for the BRI+CAS MoA, affecting mainly chitin accumulation in the cell wall. Finally, the casein kinase Yck3 is also important for the modulation of the expression of genes involved in the cell wall construction.

### BRI and BRI+CAS affect *C. neoformans* virulence

Next, we investigated if BRI+CAS affected *C. neoformans* virulence and its determinants, such as capsule formation, and melanization. It has been speculated that the inefficacy of CAS in *C. neoformans* is because the cell wall or polysaccharide capsule prevent the accessibility of CAS to β-1,3-glucan synthase^32^. We measure if BRI, CAS, and BRI+CAS could inhibit both capsule size and formation (**Figures 8a to 8c**). We found that CAS and BRI+CAS can equally inhibit more the capsule diameter size and the number of capsular cells than the wild-type strain while there are no differences between BRI and the control (**Figures 8a to 8c**). The *C. neoformans* acapsular mutants *cap59*Δ and *cap67*Δ were similarly BRI susceptible but more CAS-susceptible than the wild-type strain (**Table 1**). Checkerboard assays showed that BRI and CAS have additive effects against *C. neoformans cap59*Δ with FIC of 0.75 (**Figure 8d**). These results suggest that although BRI can potentiate more CAS activity against *C. neoformans cap59*Δ than the wild-type, CAS affected capsule formation more than BRI. Also, we did not observed changes in *C. neoformans* melanin accumulation when the cells were exposed to BRI, CAS, and BRI+CAS either at 30 or 37 °C (**Figure 8e**).

**Figure 8.**
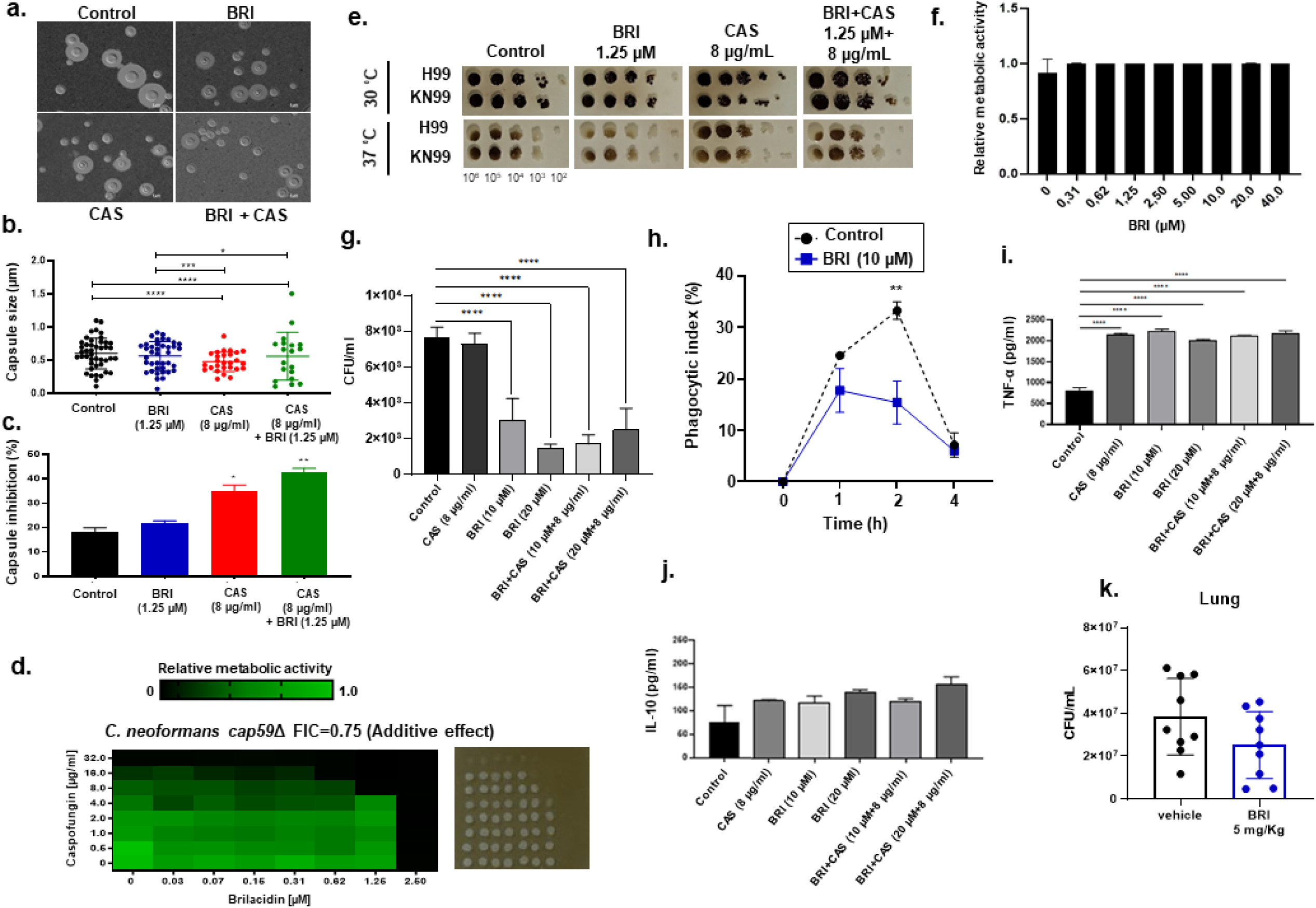
BRI and BRI+CAS affect *C. neoformans* virulence. **a.** Capsule formation after 72 h in the absence or presence of BRI (1.25 µM), CAS (8 µg/mL) or BRI+CAS (1.25 µM+8 µg/mL). **b.** Capsule thickness in the same conditions. **c.** Determination of capsule formation inhibition. **b. and c.** are based on two repetitions of 100 cells in each treatment. Statistical analysis: Ordinary one-way Anova; \**p* value<0.05; \*\*\**p* value<0.001; \*\*\*\**p* value<0.0001. **d.** Checkerboard and cidality assays using XTT. *C. neoformans cap59*Δ mutant was grown for 48 h at 30 °C with different concentrations of BRI and CAS. **e.** Melanin production was evaluated by growing *C. neoformans* H99 and KN99a strains in the absence or presence of BRI (1.25 µM), CAS (8 µg/mL), or BRI+CAS (1.25 µM+8 µg/mL). The plates were incubated for 5 days at 30 °C or 37 °C, in the dark. **f.** Bone marrow-derived macrophages (BMDMs) were exposed to various concentrations of BRI for 24 h and metabolic activity measured with XTT. **g.** *C. neoformans* viability 24 h after engulfment by BMDMs in the absence or presence of BRI (10 or 20 µM), CAS (8 µg/mL) or BRI+CAS (10 or 20 µM+8µg/mL). **h.** Phagocytic index (%) of BMDMs infected with *C. neoformans* exposed or not to 10 µM of BRI. Two repetitions of 100 cells ± standard deviation were used in each treatment. Statistical analysis: Two-way Anova Sidak’s multiple comparisons test; \*\**p* value<0.005. **i. and j.** TNF-α and IL-10 determination 24 h after BMDMs were infected or mock-infected with *C. neoformans*. **k.** *C. neoformans* lung burden 13 days after intranasal infection with 1×10^6^ cells of *C. neoformans*. Statistical analysis: one-sided Welch’s test-t, *, *p-*value: 0.05.

*C. neoformans* may be engulfed by host macrophages, potentially remaining latent inside of them or taking advantage of them for dissemination within the host^75^. We found that BRI is not toxic to bone marrow-derived macrophages (BMDMs; **Figure 8f**) and that BRI or BRI+CAS decrease *C. neoformans* survival in BMDMs about 80 to 90 % (**Figure 8g**). Considering the possibility that BRI changes the *C. neoformans* macrophage recognition due to possible changes in the capsule and cell wall previously reported, we also quantified a kinetics of *C. neoformans* phagocytosis in the presence and absence of BRI 10 µM (**Figure 8h**). In the absence of BRI, we observed phagocytic indexes of 24.6, 33.3, 7.2 %, while in the presence of BRI 10 µM, phagocytic indexes of 17.8, 15.4, and 6 %, in 1, 2, and 4 h phagocytosis, respectively (**Figure 8h**). The phagocytic index in the presence of BRI in 2 h is significantly reduced 54 % from the corresponding phagocytic time in the absence of BRI (**Figure 8h**). These results suggest that BRI can affect both the initial steps of *C. neoformans* phagocytosis and initial and later steps of viability in the presence of BMDMs.

We also examined the host response to cryptococcal cell exposure in the presence of these compounds. CAS, BRI and BRI+CAS are affecting the BMDMs TNF-α production but did not change the levels of the anti-inflammatory cytokine IL-10 during *C. neoformans* BMDM infection (**Figures 8i to 8j**). Importantly, BRI treatment BRI (5 mg/Kg, administered once daily by intraperitoneal injection for 13 days) can significantly decrease (about 34.5 %) *C. neoformans* lung infection in an immunocompetent murine model of invasive pulmonary cryptococcosis (**Figure 8k**).

Taken together, our results suggest that BRI has the potential to affect *C. neoformans* virulence and pathogenicity.

## Discussion

*C. neoformans* is an important fungal pathogen that affects a large global population of people with immunosuppressive conditions, including HIV patients. There are very few options to treat cryptococcal infections since *Cryptococcus spp.* are intrinsically resistant to echinocandins, and azoles are not very efficient because there are several mechanisms to develop azole resistance^19,22,76,77^. The primary treatment for initial therapy in disseminated or central nervous system (CNS) cryptococcosis is AmB, which may be used alone or in combination with 5-fluorocytosine. AmB has a rapid onset of action and often leads to clinical improvement more rapidly than either intravenous or oral fluconazole; after that, patients usually need to take fluconazole for an extended time to clear the infection. However, amphotericin causes substantial toxicity and its application is limited by economic issues^22^. Recently, other drugs, such as fosmanogepix (which inhibits the GPI-anchor biosynthetic enzyme Gwt1), sertraline (a repurposed antidepressant that inhibits serotonin reuptake), tamoxifen (a repurposed breast cancer therapeutic; that selectively modulates the oestrogen receptor), and VT-1598 (an inhibitor of the ergosterol biosynthetic enzyme Erg11 that disrupts membrane integrity) are at different stages of clinical trial testing and are promising novel antifungal agents to treat cryptococcosis (for a review, see ^22^). Here, we demonstrate that the host defense peptide mimetic BRI is a novel compound with toxicity for not only reference strains and clinical isolates of *C. neoformans* but also *C. gattii*. BRI is efficient by itself but also can potentiate CAS and AmB against both species. BRI may thus offer a therapeutic alternative for cryptococcal infection.

Antimicrobial peptides directly or indirectly target microbial plasma membranes, disrupting their membrane potential^78^. BRI has been shown to act by a similar mechanism in various bacterial species^54,79,80^. We have previously shown that BRI can potentiate CAS and azoles against *A. fumigatus*, *Candida albicans*, *C. auris, C. neoformans* and several species of mucorales^41^. Here, we investigated the MoA of BRI and how BRI can potentiate CAS against *C. neoformans*. By using specific mutants and chemical inhibition, we demonstrate that BRI affects the organization of the *C. neoformans* cell membrane, influencing its permeability. Ergosterol is the major sterol in fungal membranes and is critical for the establishment of membrane fluidity and the regulation of cellular processes. Glycosphingolipids (GSLs) are key components of the plasma membrane and are involved in cellular processes crucial for fungi, such as growth, differentiation, signal transduction, and pathogenesis^53^. GSLs are clustered along with sterols in specialized membrane microdomains termed lipid rafts, which are essential for cell membrane organization and crucial for the establishment of cell polarity^53,81^. The organized construction of the cell membrane, with the appropriate distribution of GSLs and ergosterol, is needed for the correct assembly of essential cell membrane proteins as well as the fusion and deposition of vesicles containing precursors required for cell wall growth that are transported to the hyphal tip through a network of microtubules and the actin cytoskeleton^53,82^. Our results suggest BRI most likely affects the *C. neoformans* cell membrane organization by disrupting membrane potential and perturbing the distribution of ergosterol and sphingolipids. This will affect the cell homeostasis, accelerating the process of cell death. BRI may also influence the anchorage of the cell wall to the plasma membrane, affecting either cell wall morphology or its adherence to the plasma membrane (for reviews, see ^83–85^). In both cases, altered concentrations of long-chain ceramides and complex sphingolipids could affect the rigidity of the cell wall and/or cell membrane (see ^86^; and for reviews, see ^83–85^).

Changes in cell membrane composition and organization also affect protein secretion, endocytosis, and the proper deposition of components of the cell wall and plasma membrane (for a review, see ^87^). Consistent with this, we found that several mutants involved in endocytic traffic were more sensitive to BRI than wild-type cells and displayed increased permeability to PI when exposed to BRI. Membrane disorganization can also influence the correct organization and positioning of receptors, membrane proteins and membrane receptors that are essential for regulation of signaling transduction pathways essential for cell survival. We used the power of *S. cerevisiae* molecular genetics to identify gene products important for the cellular response to BRI. We observed a very complex multitarget MoA for BRI against *S. cerevisiae*, revealing multiple mutants affected in ergosterol and sphingolipid biosynthesis to be highly susceptible to BRI; other mutants important for cytoskeleton and cell wall biosynthesis were depleted or enriched after drug treatment. More interestingly, mutants lacking protein kinase C, MAP kinases from the CWI and HOG pathways, and calmodulin have altered survival upon BRI treatment, strongly indicating that BRI by itself is primarily influencing these pathways.

BRI impact is particularly evident when *C. neoformans* is exposed to CAS and there is an increase of the exposure of chitin or changes in the thickness in the *C. neoformans* cell wall that are repressed either in the presence of BRI or BRI+CAS. These effects are also observed upon transcriptional regulation of genes important for the biosynthesis of different components of the cell wall. Our results indicate that BRI is hierarchically able to affect the cell membrane and cell wall organization. The observed CAS synergism could be related to an increasing worsening phenotype of cell wall disconstruction through the β-1,3-glucan depletion which will promote accentuated permeability and leakage of the cytoplasmic content and increased lethality observed upon BRI+CAS. This is further supported by our identification of casein kinase inhibitors when we screened for PKIs that could potentiate BRI. The nonessential *C. neoformans* casein kinase 1 *cck1/ypk3*^61,62^ has several deficient phenotypes related to cell membrane homeostasis and the cell wall integrity pathway, and regulates the phosphorylation of both Mpk1 and Hog1 mitogen-activated protein kinases (MAPKs) during cell wall and cell membrane stresses^61,62^. Comparatively, the *Candida albicans* casein kinase 1 family member *yck2*Δ/*yck2*Δ mutant was hypersusceptible to cell wall damaging agents, had increased chitin content in the cell wall, and increased mRNA accumulation of the chitin synthase genes, *CHS2*, *CHS3*, and *CHS8*^88^. A screening of protein kinase inhibitors to reverse *C. albicans* CAS resistance identified a compound able to restore CAS sensitivity whose target is Yck2^60^. Recently, a screening for *C. albicans* kinome to identify genes for which loss-of-function confers hypersensitivity to echinocandins and azoles revealed the casein kinase 1 (CK1) homologue Hrr25 as a regulator of tolerance to both antifungals and especially important as a target-mediated echinocandin resistance^89^. BRI potentiates the action of CAS against *C. albicans*^41^, but it remains to be determined if *C. albicans* casein kinase mutants have increased BRI susceptibility. *C. neoformans* Cck1/Ypk3 modulates the expression at transcriptional level of genes encoding ergosterol biosynthesis, β-1,3-glucan synthase, and chitin synthase. Other *C. neoformans* protein kinases that interact with casein kinases, such as calcineurin and MAP kinases, also have decreased BRI susceptibility.

When we investigated the influence of BRI on several key *C. neoformans* virulence determinants, we found that BRI decreases capsule formation but not melanin production. BRI also significantly reduces *C. neoformans* phagocytosis and survival inside macrophages, despite being nontoxic to BMDMs. It partially clears *C. neoformans* lung infection in an immunocompetent murine model of invasive pulmonary cryptococcosis. Interestingly, CAS and BRI can increase the TNF-α but not the IL-10 cytokines production. TNF-α plays a critical role in the control of *C. neoformans* and its lack stimulates its persistence^90^ while IL-10 signaling has been shown as important for persistent cryptococcal lung infection^91^. These results indicate that BRI is able to immunomodulate cytokine production and decrease the *C. neoformans* infection.

We have demonstrated that BRI is potentially an important antifungal agent against cryptococcosis, particularly important for synergizing with CAS, which is otherwise ineffective against this pathogen. Future work will determine how BRI interacts physically with components of the cell membrane, and how this affects CWI.

## Methods

### Ethical statement

The principles that guide our studies are based on the Declaration of Animal Rights ratified by UNESCO on January 27, 1978 in its 8th and 14^th^ articles. All protocols adopted in this study were approved by the local ethics committee for animal experiments from the University of São Paulo, Campus of Ribeirão Preto, Brazil (Permit Number: 08.1.1277.53.6; Studies on the interaction of fungal pathogens with animals).

### Strains, media and cultivation methods

All the *Cryptococcus* strains are shown in **Table 1**. For all studies, *C. neoformans* and *C. gattii* strains were inoculated from single colonies into YPD media (2% [wt/vol] dextrose, 2% [wt/vol] Bacto peptone, and 1% [wt/vol] yeast extract in distilled water [dH_2_O]) and grown overnight at 30°C with shaking at 160 rpm before further handling as detailed below. Agar was added to a final concentration of 1.5% when solid medium was used. RPMI-1640 media (Gibco) supplemented with 9.6 mM HEPES (Sigma-aldrich) was used for metabolic activity assays and MIC assays.

### Minimal inhibitory concentration (MIC)

The BRI drug used for MIC assays was kindly supplied by Innovation Pharmaceuticals Inc. and solubilized in DMSO. The minimal inhibitory concentration (MIC) for *Cryptococcus strains was* determined based on the European Committee on Antimicrobial Susceptibility Testing (EUCAST)^92^, using E246-7 method. In brief, the MIC assay was performed in 96-well flat-bottom polystyrene microplates where 100 µL of a 10^5^ cells/mL stock prepared in liquid RPMI-1640 media was dispensed in each well and supplemented with increasing concentration of drugs. Plates were incubated at 30 °C for 48 h and the inhibition of growth was evaluated. The MIC was defined as the lowest drug concentration that yielded 100% fungal growth inhibition, assessed visually, compared with control wells containing only RPMI-1640 medium and DMSO.

### Culture conditions and ergosterol extraction

The extraction and quantification of ergosterol were carried out as described previously with some modifications^93^. *Cryptococcus* strains (KN99a and *erg3*△) were grown in liquid YPD medium at 30°C with shaking for 16 h. Next, cells were collected by centrifuge (3000 rpm for 10min) and washed with sterile distilled water. Cells were counted by hematocytometer and the inoculum adjusted to 10^7^ to 10^8^ cells into liquid RPMI-1640 medium with (2.5 µM or 25 µM) or without brilicidin. The cells were incubated at 30°C for 4 h and then collected and washed as above. Three milliliters of 25% ethanolic of KOH solution were added to each pellet and vortexed for 1 min. The cell suspension was transferred to borosilicate glass screwcap tubes to incubated at 85 °C for 1 h in a water bath. Sterols were extracted by adding a mixture of 1 ml of sterile distilled water and 3 mL of petroleum ether. The petroleum ether layer was transferred to a clean borosilicate glass and sterols were detected by reading absorbance at 260 nm and 280 nm.

### Checkerboard assays

For measuring the effect of the combination between BRI and antifungal drugs against *Cryptococcus* spp., metabolic activity by XTT-assay was used ^94^. A concentration of 10^5^ cells/mL were inoculated in liquid RPMI-1640 supplemented with increase concentrations of BRI (x-axis) and increase concentrations of CAS, 5-Fluorocytosine (5-FC), FLC, Latrunculin-B or AmB (y-axis) in 96-well flat-bottom polystyrene microplate. The plates were incubated at 30°C for 48 h. The cells viability were revealed using XTT-assay as described^94^. Before XTT-assay, 3 µL from each well were collected and plated on solid YPD for cidality evaluation. The plates were incubated at 30 °C for 24 h and photographed. To determine synergy, additive, indifference, or antagonism we used the FIC index method^95^; where FIC index of <0.5 indicates synergism, 0.5–4 indicates additive or indifference, and >4 is considered to be antagonism^96^.

For *S. cerevisiae*, the interaction between BRI (0-80µM) and fluconazole (0-16µg/mL), voriconazole (0-4µg/mL), CAS (0-1µg/mL) or anidulafungin (0-2µg/mL) was assessed through a checkerboard assay using liquid YPGal medium. Briefly, single colonies of cells were inoculated from solid YPD in liquid YPD and incubated for 16 h at 30°C. Cells were washed twice in liquid YPGal and ressuspended to a concentration of 2.5 x 10^3^ cells/mL in the same medium supplemented with increasing concentrations of BRI (x-axis) and increasing concentrations of fluconazole, voriconazole, CAS and anidulafungin in 96-well flat-bottom polystyrene microplate. The plates were incubated at 30°C for 48 h and the metabolic activity was measured using XTT-assay as described^94^. Before XTT-assay, the cidality was evaluated by plating 3 µL from each well on solid YPD. The plates were incubated at 30 °C for 24 h and photographed. We used the SynergyFinder2.0 (https://synergyfinder.fimm.fi) to calculate the interaction score between BRI and the other antifungal compounds. The interaction between the drugs was classified based on the synergy score where values lower than −10 were considered antagonistic, values from −10 to 10 were considered additive and values higher than 10 were considered synergistic.

### Fluorescence microscopy

For fluorescence microscopy analysis, *C. neoformans* strains were grown as above. To evaluate the relationship between BRI and CAS, *C. neoformans* H99 strain was incubated under control conditions (RPMI-1640 only) or treatments with BRI (1.25 µM), fluorescence CAS probe (CAS-F) (8 µg/mL) or with combination CAS-F+BRI ([8 µg/mL] and [1.25 µM], respectively) for 1 h at 30 °C. After the incubation time, 10 µg/mL CellTracker Blue^TM^ CMAC Dye (Invitrogen) were add into each well and incubated for 10 min in room temperature (RT). The cells were collected and washed tree times with 1x PBS (140 mM NaCl, 2 mM KCl, 10 mM NaHPO4, 1.8 mM KH_2_PO_4_, pH 7.4).

For evaluation of membrane integrity, the H99 and KN99a strains and the mutants *vps3*Δ and *erg3*Δ were incubated in liquid RPMI-1640 media under control conditions or treatment with BRI (1.25, 2.5, 12.5 and 25 µM), CAS (8 µg/mL) or the combination of BRI + CAS and incubated at 30°C for 1 h, 2h or 4 h. After, the cells were washed 2 times with 1x PBS and staining in 1x PBS with Propidium Iodide (10 µg/mL 15 min, RT) or Filipin (10 µg/mL 20 min, RT). The cells were collected and washed tree times with 1x PBS.

Slides were visualized on an Observer Z1 fluorescence microscope using a 100x objective oil immersion lens for Filipin, filter set 38-high efficiency [HE], excitation wavelength of 450 to 490 nm, and emission wavelength of 500 to 550 nm; and for PI, the wavelength excitation was 572/25 nm and emission wavelength was 629/62 nm, Filter Set 63 HE. DIC (differential interference contrast) images and fluorescent images were captured with an AxioCam camera (Carl Zeiss) and processed using AxioVision software (version 4.8). In each experiment, at least 50 cells were observed and the experiment repeated at least two times.

### *S. cerevisiae* chemical genomics analysis

Chemical genomics analysis using *S. cerevisiae* mutant libraries was conducted as described^58,97^. The libraries include temperature-sensitive (TS), overexpression (MoBY), and diploid heterozygous deletion (HET) collections. A haploid deletion collection (ScWG: *<u>S</u>accharomyces <u>c</u>erevisiae* <u>w</u>hole <u>g</u>enome), which is a barcoded collection of ∼3,500 haploid non-essential gene deletions in the drug hypersensitive strain (Y13206: *MATα snq2Δ:: KlLEU2 pdr3Δ:: KlURA3 pdr1Δ:: NATMX can1Δna11:: 2iSp_his5 lyp1Δ his3Δ1 leu2Δ0 ura3Δ0 met15Δ LYS2*), was also constructed using SGA technology (manuscript in preparation) and used for chemical genomics analysis. Pooled yeast mutant libraries were treated with BRI. Basically, the workflow including culture, DNA extraction, and PCR amplification of each strain-specific barcode proceeded as described in ^58^. Purified PCR products were sequenced with an Illumina Miseq machine at the RIKEN Center for Brain Science (Wako, Japan).

### Transmission electron microscopy

*C. neoformans* strain KN99a was grown overnight in YPD as above, washed in distilled water, resuspended at 10^8^/mL in RPMI-1640 medium (Gibco^TM^), and dispensed (1 mL aliquots) into 24-well plates. Caspofungin diacetate and brilacidin tetrahydrochloride were added to achieve the indicated final concentrations and the plates were incubated (8 h, 30°C) before cell fixation with 2% paraformaldehyde/2.5% glutaraldehyde (Ted Pella Inc., Redding, CA) in 0.1 M sodium cacodylate buffer (2 h at room temperature and then overnight at 4°C). Samples were next washed in sodium cacodylate buffer and postfixed in 1% osmium tetroxide (Ted Pella Inc.) for 1 h at room temperature. After three washes in distilled water, samples were en bloc stained in 1% aqueous uranyl acetate (Electron Microscopy Sciences, Hatfield, PA) for 1 h, rinsed in distilled water, dehydrated in a graded series of ethanol, and embedded in Eponate 12^TM^ resin (Ted Pella Inc). Ultrathin sections of 95 nm were cut with a Leica Ultracut UCT ultramicrotome (Leica Microsystems, Bannockburn, IL), stained with uranyl acetate and lead citrate, and viewed on a JEOL 1200 EX transmission electron microscope (JEOL USA Inc., Peabody, MA) equipped with an AMT 8megapixel digital camera and AMT Image Capture Engine V602 software (Advanced Microscopy Techniques, Woburn, MA).

### Staining for cell surface componentes

Cell wall surface polysaccharide staining was performed using at least 6 repetitions. Briefly, the cells were grown overnight in YPD. Then, washed with PBS and 100 µL of 10^6^ cells/mL suspension was inoculated in RPMI-1640 medium with BRI (25 µM) and/or CAS (8 µg/mL), in 96-well microplate. The plate was incubated for 8 h at 30 °C. The culture medium was removed and the cells were fixation with PBS 3.7% formaldehyde/2.5% glutaraldehyde for 5 min at RT. For chitin staining, 100µl of a PBS with 25 µg/mL of calcofluor white (CFW) was added to the wells, which were incubated for 10 min at RT and washed twice with PBS before fluorescence was read at 380-nm excitation and 450-nm emission. The fluorescence was read in a microtiter plate reader (SynergyHTX Multimode Reader; Agilent Biotek or EnSpire Multimode Plate Reader; Perkin Elmer).

### RNA isolation, cDNA synthesis and RTqPCR analysis

All experiments were carried out in biological triplicates, *C. neoformans* H99 and *yck3Δ* strains were grown overnight in YPD as above. Cells were washed in PBS and resuspended at 10^6^ cells/mL in RPMI-1640 medium. BRI (25 µM) and CAS (8 µg/mL) were added and the inoculum was incubated for 4 h at 30 °C, 180 RPM. The cells were washed in PBS and the samples was lyophilized. A total RNA was isolated by TRIzol (Invitrogen) after cellular lysis by glass beads and treated with RQ1 RNase-free DNase I (Promega). RNA integrity and concentration were assessed using a NanoDrop Lite Spectrophotometer (Thermo Scientific). For RT-qPCR, the RNA was reverse transcribed to cDNA using the ImProm-II reverse transcription system (Promega) according to manufacturer’s instructions, and the synthesized cDNA was used for real-time analysis using the SYBR green PCR master mix kit (Applied Biosystems) in the ABI 7500 Fast real-time PCR system (Applied Biosystems, Foster City, CA, USA). Sybr Primer sequences are listed in **Supplementary Table S6 (10.6084/m9.figshare.25239550)**. *ACT1* (Actin-1) gene was used as normalizer.

### Transcription factor and protein kinase inhibitor screening

To identify possible genes involved in the mechanism of BRI action against *Cryptococcus* spp., we screened a library of *C. neoformans* H99 mutants for transcription factors. For screening the library, the frozen cells were inoculated in 600 μL liquid YPD in a deep-well plate for 48 h at 30°C, 180 rpm. After incubation, 100 µL of each well were transferred to a clear 96-flat bottom-plate and optical densities measured at 600 nm; cells were normalized to an OD_600_ of 1 in a final volume of 100 μL YPD. Then, cells were diluted 1:20 in 200 μL liquid RPMI-1640 and adjusted to a final concentration of 10^5^ cells/mL in liquid RPMI-1640 media supplemented with BRI (1.25 µM) or CAS (8 µg/mL), in 2 replicate plates. Fresh liquid RPMI-1640 with DMSO was used as control. The plates were incubated at 30°C for 48 h. The absorbance was read at 600 nm to determine the optical density and 3 µL from each well was plated on solid YPD to evaluate cell viability.

The candidates initially selected as sensitive to BRI or CAS were grown overnight in liquid YPD at 30°C, 180 rpm. Then, cells were washed twice with 1x PBS and cellular density was adjusted to 10^6^ cells/mL. A microdilution was performed and the cells were grown in a 96-flat bottom-plate on liquid RPMI-1640 media in the presence (or not) of BRI (1.25 µM) or CAS (8 µg/mL) for 48 h at 30°C. After that, 3 µL from each well was plated on solid YPD and incubated at 30°C for 24 h. Cell viability was assessed visually and photographed.

A protein kinase inhibitor (PKI) library was also screened in combination with BRI. In total, 58 PKI were analyzed (**Supplementary Table S5; 10.6084/m9.figshare.25239550**). Briefly, 100 µL of liquid RPMI-1640 containing 10^5^ cells/mL of the *C. neoformans* H99 strain was inoculated with PKI (20, 40 and 60 µM) in the presence (or not) of BRI (1.25 µM) and incubated 48 h at 30°C. The cells were then collected and plated on solid YPD for CFU counting. Experiments were repeated at least two times.

### Luria-Delbruck fluctuation assay

Fluctuation assays were conducted as previously described^98^. Briefly, ten independent H99 colonies were selected from a YPD agar stock and cultured overnight in 5 mL liquid YPD at 30°C. The cultures were washed 2x with dH_2_O, resuspended in 5 mL dH_2_O, and plated onto the appropriated solid RPMI medium (100 μL of cells at a 10^−5^ dilution were plated onto RPMI, and 100 μL undiluted cells onto RPMI + 5-FC [100 µg/mL] and RPMI-1640 + BRI [100 µM]; no repeated measurements). The plates were incubated at 37°C; colonies were counted following incubation for 4 (control RPMI-1640 medium) or 14 days (drug media). Data from the fluctuation assay was analysed using the R package flan v0.9^99^. The analysis was conducted with default parameters, employing a 95 percent confidence interval for the mutation probability and the maximum likelihood method.

### Capsule formation analysis

To qualitatively assess capsule thickness, H99 strain were grown on YPD medium for 16 h and washed with PBS, and 10^5^ cells were incubated in Dulbecco’s MEM media (Gibco) with 10% FBS, in a control condition and in the presence of BRI (1.25 µM) or CAS (8 µg/mL), and the combination of BRI + CAS (1.25 µM and 8 µg/mL, respectively) for 72 h at 37°C and 5% CO_2_. Cells were collected and counter-stained with India ink (1:1) for microscopy analysis. Relative capsule sizes were defined as the ratio between the capsule thickness and cell diameter. ImageJ software was utilized to determine the capsule measurements of 50 cells of two technical replicates.

### Melanin induction

Melanin production was evaluated by growth of *C. neoformans* H99 and KN99a strains in a solid minimal medium (MM [15 mM glucose, 10 mM MgSO_4_, 29.4 mM KH_2_PO_4_, 13 mM glycine, 3 μM thiamine, pH 5.5]), with 1 mM L-DOPA (Sigma-Aldrich), in a control condition and with added BRI [1.25 µM], or CAS [8 µg/mL], or BRI+CAS [1.25 µM+8 µg/mL]. The plates were incubated for 5 days at 30°C or 37°C, in the dark as described ^100^ with some adaptations.

### Macrophage culture

BALB/c bone marrow-derived macrophages (BMDMs) were obtained as previously described^101^. Briefly, bone marrow cells were cultured for 7–9 days in Dulbecco’s Modified Eagle Medium (DMEM) 20/30, which consists of DMEN medium (Gibco, Thermo Fisher Scientific Inc.), supplemented with 20 % (vol/ vol) FBS and 30 % (vol/vol) L-Cell Conditioned Media (LCCM) as a source of macrophage colony-stimulating factor (M-CSF) on non-treated Petri dishes (Optilux - Costar, Corning Inc. Corning, NY). Twenty-four h before experiments, BMDMs were detached using cold phosphate-buffered saline (PBS) (Hyclone, GE Healthcare Inc. South Logan, UT), the cell viability and concentration was adjusted using a hemacytometer as previously described^102^ diluting 1 part of 0.4% trypan blue (Gibco) and 1 part cell suspension. After count the unstained (viable) and stained (nonviable) cells separately, the percentage of viable cells was calculated as follows:

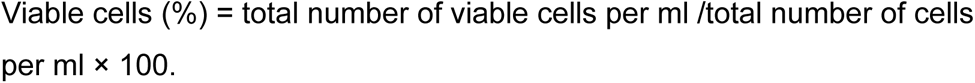

For functional assays 1 × 10 cells were added to 24-well tissue culture plates containing the differentiation Eagle *Medium* (*DMEM)* 20/30 and allowed to grow adherently overnight at 37^◦^C. Non-adherent cells were removed by gently washing three times with warm PBS. The viability of the BMDMs preparations was >99% as judged by trypan blue dye exclusion and the cells percentage is greater than 90% differentiated macrophages.

### Macrophage internalization and killing assay

Bone marrow-derived macrophages were prepared as described above. Briefly, 10^6^ cells/well wereseeding in a 24-well plate and incubating in DMEM supplemented with 10% fetal bovine serum (FBS) at 37°C and 5% CO_2_ for 24 h. *C. neoformans* cells were prepared for uptake experiments by inoculating an overnight culture in YPD and growing at 30°C, 180 rpm. To initiate the study, cryptococcal cells were washed with PBS and opsonized with anti-glucuronoxylomannan antibody 18B7 (1 μg/mL, Sigma)for 1 h at 37°C, while macrophages were activated with 1 µg/mL liposaccharide (LPS) for 30 min at 37°C and 5% CO_2_; 10^7^ cryptococcal cells were then incubated with the macrophages at 37°C and 5% CO_2_. After 24 h, the wells were washed three times with warm 1x PBS, and the macrophages were lysed with dH_2_O with 0.1% Triton X-100. This suspension was then diluted and 25 μL was plated on solid YPD and incubated at 30 °C for 24 h for CFU counting. Also, the cell supernatant was collected and reserved for cytokine quantification assays.

### Phagocytosis assay

Bone marrow-derived macrophages were prepared as described above. Briefly, 5 x 10^5^ cells/well were seeding in a 24-well plate containing circular glass coverslips and incubating in DMEM supplemented with 10% fetal bovine serum (FBS) at 37°C and 5% CO_2_ for 24 h. *C. neoformans* cells were prepared for uptake experiments by inoculating an overnight culture in YPD and growing at 30°C, 180 rpm. To initiate the study, cryptococcal cells were washed with PBS and opsonized with anti-glucuronoxylomannan antibody 18B7 (1 μg/mL, Sigma) for 1 h at 37°C. After the incubation, the cells were washed and incubated with 0.1 mg/ml FITC (Sigma) in 0.1 M Na_2_CO_3_ at 37 °C for 30 min. While macrophages were activated with 1 µg/mL liposaccharide (LPS) for 30 min at 37°C and 5% CO_2_; Labelled cells were washed three times with PBS, and 5 x 10^6^ cryptococcal cells were then incubated with the macrophages at 37°C and 5% CO_2_. After 2 h, labelling of extracellular cryptococcal cells was performed by incubation with PBS, 0.25 mg/ml calcofluor white (Sigma) for 30 min at 4°C. The cells were washed twice with PBS and fixed with 3.7 % (vol/vol) formaldehyde/PBS for 15 min followed by two washes with PBS. Microscopic photographs were taken on a Zeiss microscope.

### Animal procedures

Inbred female mice (BALB/c strain; body weight, 20–22 g; obtained from the facility of the campus of the University of São Paulo, Ribeirão Preto, Brazil) were used in this study. Mice were housed 5 per cage and had access to food and water *ad libitum*. Cages were well ventilated, softly lit and subjected to 12:12 h light-dark cycle. The relative humidity was kept at 40 – 60% and mouse room cages were kept at 22°C. The *C. neoformans* H99 strain was sub-cultured at 30°C for 48 hours on solid YPD medium. Prior to inoculation, colonies were taken from the subculture and inoculated into liquid YPD and grown overnight at 30°C, 180 rpm; cells were collected by centrifugation and washed three times in PBS. Mice (10 mice per group) were anesthetized by halothane inhalation and infected by intranasal instillation of 20 µL 10^6^ cells of *C. neoformans* H99. The viability of the administered inoculum was determined by incubating a serial dilution of the cells in solid YPD, at 30°C. Treatment groups will consist of a vehicle (Cavitron^TM^ W7 HP7 from Ashland, day 1 - 13) and BRI (10 mg/Kg, day 1; 5 mg/Kg, day 2 - 13). The drugs used in the mice treatement were prepared on day 1 and again on day 7, and stock refrigerated. For vehicle solution preparation, 20% Cavitron^TM^ W7 HP7 was diluted in sterile water for injection and filtered in 0.22 µm sterile filters. For brilacidin treatment, stock dosing solutions was prepared from the received brilacidin tetrahydrochloride dry powder (Innovation Pharmaceuticals), diluted in vehicle solution and filtered in 0.22 µm sterile filters. The drugs were prepared to deliver their respective dose within a 0.1 mL volume. Animals (10 per group) were treated for 13 days (started in the day 1 post-infection) once daily by intraperitoneal (IP) injection. Mice were sacrificed 14 days post-infection. Animals were clinically monitored at least twice daily. Any animal that appears moribund prior to the scheduled endpoint was humanely euthanized. As a negative control, a group of 10 mice received vehicle only. To investigate fungal burden, the lungs were harvested and stored in ice. Samples were homogenized with sterile PBS with protease inhibitor cocktail tablets complete Tablets, mini EDTA-free (Roche) and homogenates adequately diluted for fungal burden evaluation by colony forming units.

### Cytokine quantification

ELISA-assay was used to quantify the cytokine levels in cells supernatant, after 24 h of exposure to *C. neoformans* H99. The quantification of cytokines Tumor Necrosis Factor (TNF-α) and Interleukin-10 (IL-10) was performed according to the manufacturer’s instructions, using ELISA-assay kits (R&D Systems). The plates final absorbance was read at 450 nm and the cytokine concentration analysis [pg/mL] was performed according to the manufacturer’s instructions, considering the values obtained in the standard curve of each evaluated cytokine.

### Statistical analysis

Grouped column plots with standard deviation error bars were used for representations of data. For comparisons with data from wild-type or control conditions, we performed one-tailed, paired *t tests* or one-way analysis of variance (ANOVA). All statistical analyses and graphics building were performed by using GraphPad Prism 8 (GraphPad Prism Software).

## Data availability statement

The datasets generated for this study are available on request to the corresponding author.

## Acknowledgements

We thank the Fundação de Amparo à Pesquisa do Estado de São Paulo (FAPESP) grant numbers 2021/04977-5 (GHG) and 2022/08796-8 (CD), 2021/10599-3 (The Antimicrobial Resistance Institute of São Paulo, The Aries Project) and the Conselho Nacional de Desenvolvimento Científico e Tecnológico (CNPq) grant numbers 301058/2019-9, 404735/2018-5, and 405934/2022-0 (The National Institute of Science and Technology INCT Funvir) (GHG), both from Brazil, the National Institutes of Health/National Institute of Allergy and Infectious Diseases from the USA, grants R01AI153356 to GHG, R01AI053721 to JWK, and R21AI178330 to TLD, JSPS KAKENHI numbers 17H06411 (YY and CB) and 23H04882 (YY) from Japan. This work was funded by the Joint Canada-Israel Health Research Program, jointly supported by the Azrieli Foundation, Canada’s International Development Research Centre, Canadian Institutes of Health Research, and the Israel Science Foundation (GHG). We also thank David Harold Drewry for providing the protein kinase inhibitor library, and RIKEN Center for Life Science Technologies and the Support Unit for Bio-Material Analysis for sequencing. We also thank Michael Bottery for his suggestions about the Luria-Delbruck fluctuation test, Thiago Aparecido da Silva and Andrew Alspaugh for providing *Cryptococcus spp* strains, and Jane Harness for critical Reading of the manuscript. CB and JWK are Fellows in the CIFAR Fungal Kingdom: Threats & Opportunities program. CB was supported by Canadian Institutes of Health Research (FDN-143264).

## Author Contributions Statement

CD, CFP, PAC, ED, LGC, ESL, KB, DgK, SA, NNM, MY, LAVI, LTKP, YY, and TFR performed most of the experiments. JWK, TLD, ASI, and CB provided advices and materials for the work. GHG and CD analyzed the data, wrote the manuscript and GHG coordinated all the work. All the authors read and edited the manuscript.

## Supplementary Figures and Tables

**Supplementary Figure S1.** Bokeh plots for the mutants depleted or enriched in *S. cerevisiae* deletion library.

**Supplementary Figure S2.** Bokeh plots for the mutants depleted or enriched in *S. cerevisiae* haploinsuficiency library.

**Supplementary Figure S3.** Bokeh plots for the mutants depleted or enriched in *S. cerevisiae* overexpression library.

**Supplementary Figure S4.** Bokeh plots for the mutants depleted or enriched in *S. cerevisiae* temperature-sensitive library.

**SupplementartyFigure S5.** Checkerboard and cidality assays for *C. neoformans* grown for 48 hs at 30 °C

**Supplementary Table S1.** Genes observed as depleted or enriched when the deletion library is grown in the presence of different brilacidin concentrations.

**Supplementary Table S2.** Genes observed as depleted or enriched when the haploinsuficiency library is grown in the presence of different brilacidin concentrations.

**Supplementary Table S3.** Genes observed as depleted or enriched when the overexpression library is grown in the presence of different brilacidin concentrations.

**Supplementary Table S4.** Genes observed as depleted or enriched when the temperature sensitive library is grown in the presence of different brilacidin concentrations.

**Supplementary Table S5.** Protein kinase inhibitors analysed in this study

**Supplementary Table S6.** Primers used in this work.

